# *Medicago truncatula* SOBIR1 controls specificity in the Rhizobium-legume symbiosis

**DOI:** 10.1101/2023.10.15.561875

**Authors:** Baptiste Sarrette, Thi-Bich Luu, Alexander Johansson, Judith Fliegmann, Cécile Pouzet, Aurélie Le Ru, Carole Pichereaux, Céline Remblière, Laurent Sauviac, Noémie Carles, Emilie Amblard, Valentin Guyot, Maxime Bonhomme, Julie Cullimore, Clare Gough, Christophe Jacquet, Nicolas Pauly

## Abstract

*Medicago truncatula* Nod Factor Perception (MtNFP) is a lysin-domain Receptor-Like Kinase (LysM-RLK) that plays a key role in the Rhizobium-legume symbiosis, and is involved in plant immunity. MtNFP also has an inactive kinase domain, suggesting that the protein is involved in different receptor complexes. Using the MtNFP pseudo-kinase domain as a bait in a Yeast two Hybrid screen, we identified *M. truncatula* SUPPRESSOR OF BIR1 (MtSOBIR1) as a new interactor of MtNFP. We showed that an interaction between the two RLKs can occur *in planta* and that the kinase domain of MtSOBIR1 is active and can transphosphorylate the pseudo-kinase domain of MtNFP. Like in other plants, our data suggest a positive role of MtSOBIR1 in immunity; *MtSOBIR1* could functionally complement an *Atsobir1* mutant for defence activation, and a *Mtsobir1* mutant was defective in pathogen-induced defence gene expression. We also showed that *MtSOBIR1* has a symbiotic role with *Mtsobir1* mutants showing a strong symbiotic phenotype in a plant genotype- and rhizobial strain-specific manner. The symbiotic role was apparent both at an early stage of rhizobial infection and in nodules. Together, these data suggest that, like MtNFP, MtSOBIR1 has a dual role, and can control immunity in both pathogenic and beneficial situations, with positive or negative roles, respectively.

## Introduction

The Rhizobium-Legume (RL) symbiosis is an intimate interaction that results in huge benefits to host plants in terms of nitrogen nutrition. Indeed, legume crops can be grown without the addition of nitrogen fertiliser, making them important components of sustainable agriculture. A nitrogen source in the form of ammonium is provided to plants thanks to the nitrogenase activity of rhizobial bacteria that are present in massive numbers within nodules, usually formed on roots of host plants. Using model legumes like *Medicago truncatula* and *Lotus japonicus*, the molecular genetic mechanisms of nodule formation and nitrogen fixation have been extensively dissected. Consequently, many key symbiotic components have been identified and characterised both in rhizobia and in plants. Many legume host plants require their rhizobial partner to produce LipoChitoOligosaccharide (LCO) signalling molecules called Nod factors. In these cases, the specific recognition by plants of Nod factors activates signalling pathways that lead to massive transcriptional changes responsible for two parallel, but coordinated, symbiotic processes; rhizobial infection and nodule organogenesis.

One of the key plant genes that controls the perception of Nod factors is called *Nod Factor Perception, or MtNFP* in *M. truncatula*, and *Nod Factor Receptor kinase 5, or LjNFR5*, in *L. japonicus* (Singh and Verma 2023). In loss of function *Mtnfp* and *Ljnfr5* mutants, no Nod factor responses are detected and nodulation is completely absent. MtNFP/LjNFR5 is a plasma membrane located lysin motif domain Receptor-Like Kinase (LysM-RLK), with three extracellular LysM domains, and is implicated in recognising Nod factors. The intracellular domain of MtNFP/LjNFR5 has no detectable autophosphorylation activity and is considered a pseudo kinase (Arrighi et al. 2006). It is likely that MtNFP/LjNFR5 associates with one or more other protein to intracellularly transmit the extracellular detection of Nod factors. In *M. truncatula*, MtNFP interacts with MtLYK3, another LysM-RLK that is essential for nodulation (Moling et al. 2014). The inactive kinase domain of MtNFP can be weakly transphosphorylated by the active kinase of MtLYK3 (Fliegmann et al. 2016). *MtLYK3* is part of a cluster of closely-related LysM-RLKs (Limpens et al. 2003; Luu et al. 2023) and other genes in this cluster may also interact with NFP and play roles in nodulation. For example, in the *M. truncatula* genotype R108, MtLYK2bis was recently shown to be another MtNFP partner, and involved in extending nodulation ability to certain natural strains of rhizobia as well as to a *Sinorhizobium meliloti* 2011 rhizobial mutant producing modified Nod factors (Luu et al. 2022).

Another interesting aspect of MtNFP is its involvement in plant immunity. *Mtnfp* loss of function mutants are more susceptible to the oomycete and fungal plant pathogens *Aphanomyces euteiches*, *Colletotricum trifolii*, *Phytophthera palmivora* and *Verticillium albo-atrum* (Rey et al. 2013) (Rey et al. 2015) (Ben et al. 2013). This function of MtNFP is considered to be independent of Nod factors, since at least in the case of *A. euteiches*, sensitive methods did not detect any LCO production by this oomycete (Rush et al. 2020). MtNFP apparently has an additional, Nod factor dependent function in immunity, as pre-incubation of plants with Nod factors reduces the induction of reactive oxygen species by pathogen components, like exudates of *A. euteiches* and chitin oligomers, a response that is defective in *Mtnfp* mutant plants (Rey et al. 2019) (Feng et al. 2019). This may be interpreted as a role of NFP in reducing plant defences to Nod factor-producing rhizobia in order to allow rhizobial infection. The necrotic phenotypes of *M. truncatula* mutants in defence regulating genes (Bourcy et al. 2013); (Berrabah, Bourcy, Eschstruth, et al. 2014); (Wang, Yu, et al. 2016); (Sinharoy et al. 2013) show the importance of the tight regulation of plant immunity for a successful symbiotic interaction, and therefore it is likely that one or both of these immunity functions of MtNFP is relevant for symbiosis. However, little is known about these functions, but they probably require interaction with additional components.

SUPPRESSOR OF BIR1 (SOBIR1) is a leucine-rich repeat (LRR) receptor-like kinase (RLK), classified as a protein kinase of the RLK-Pelle-LRR-XI-2 family (according to the iTAK classification (Zheng et al. 2016). SOBIR1 was initially discovered as the counter-player of BAK1-INTERACTING RLK1 (BIR1), a negative regulator of immunity and cell-death in *Arabidopsis thaliana (Gao et al. 2009)*. The *bir1-1* mutant shows constitutive cell-death and defence responses, which can be partially suppressed in the double mutant *bir1/sobir1*. Follow up analysis, showed that an overexpression line of *SOBIR1* exhibited constitutive immune responses including cell death and upregulation of PR genes, confirming a role of *SOBIR1* as a positive regulator of immunity (Gao *et al*., 2009), as also demonstrated in *sobir1* mutants that are more susceptible to fungal pathogens (Zhang et al. 2013); (Liebrand, van den Burg, and Joosten 2014). In another study, *SOBIR1* was found to control the inhibition of shedding of floral organs, by promoting the internalization and recycling of receptor complexes at the plasma membrane, and was designated as *EVERSHED (EVR)* (Leslie et al. 2010). SOBIR1/EVR has an active kinase domain that can auto-phosphorylate on Serine/Threonine and Tyrosine residues *in vitro* (Leslie *et al*., 2010) (Mitra et al. 2015) (Wei et al. 2022). Further studies on SOBIR1, including in other plants, have shown that it is a common coreceptor for many different LRR - receptor-like proteins (RLPs) which lack an intracellular signalling domain (Liebrand *et al*., 2014; (Snoeck, Garcia, and Steinbrenner 2023). SOBIR1 can also interact in tripartite complexes that include the BRASSINOSTEROID INSENSITIVE1-ASSOCIATED RECEPTOR KINASE1 (BAK1) (Albert et al. 2015)((van der Burgh et al. 2019).

In order to identify new partners of MtNFP with roles in symbiosis and/or immunity, we screened a yeast two-hybrid cDNA library using the KD of MtNFP as bait. Here, we describe the identification and characterisation of a new interactor of MtNFP, a protein we have called MtSOBIR1 based on homology and functional analysis. Given that SOBIR1 acts in receptor complexes as a positive regulator of immunity in many plants, we have addressed the possible function of MtSOBIR1 in immunity against pathogens and in symbiosis.

## Materials and methods

### Bacterial strains

WT strains of *Sinorhizobium meliloti* 2011 and *Sinorhizobium medicae* WSM419 carrying lacZ reporter genes were used. For *S. meliloti* 2011, the previously constructed strain *S. meliloti* 2011 (pXLGD4) was used (Luu et al. 2022). For *S. medicae* WSM419, the plasmid pLS310-1 was constructed by subcloning the *BamH*I-*EcoR*I fragment containing the full length *lacZ* coding sequence from pQF-lacZ (Kaczmarczyk et al., 2013) into the similarly cut pCM62 plasmid (Marx and Lidstrom, 2001). Competent cells of WSM419 were then electro-transformed with pLS310-1 DNA as described (Ferri et al. 2010). Both strains were grown on TY medium supplemented with tetracycline (10 µg/ml).

### Yeast two hybrid screening

A yeast two hybrid screen was performed as a custom service using the MtNFP kinase domain (aa 273-595) as bait. The corresponding DNA sequence was PCR amplified and cloned into pB27, as a C-terminal fusion to LexA (NLexA-NFP-C). The cDNA library used as prey was derived from *M. truncatula* A17 roots non-inoculated and inoculated with *A. euteiches* (Hybrigenics Services, Evry, France). The library was constructed with mRNA from a mixture of 2-week-old *in vitro* grown, non-inoculated plants, and roots from the same age plants but inoculated with *A. euteiches* and harvested at 1 and 6 days after inoculation, according to a 2/1/1 ratio. It is available as “Whole plant and root, *A. euteiches* infected, ref: [MTA17]” (Hybrigenics services). Hybrigenics Services performed their optimized ULTImate Y2H™ technique and provided information to separate artefacts from specific interactions by the global predicted biological score (PBS), which is based on a statistical model (Rain et al., 2001). The LexA bait construct was used to screen 62.5 million clones using a mating approach, with HGX13 (Y187 *ade2-101::loxP-kanMX-loxP, MATα_)* and L40_Gal4 (*MATa*) yeast strains as previously described (Fromont-Racine et al, 1997). A total of 355 His+ colonies were selected on a medium lacking tryptophan, leucine and histidine. The prey fragments of positive clones were amplified by PCR and sequenced at their 5’ and 3’ junctions. The resulting sequences were used to identify the corresponding interacting proteins in the GenBank database (NCBI) using a fully automated procedure. Following sequence analyses of these clones, a confidence score (Predicted Biological Score-PBS) was attributed to each interaction and a total of 53 genes were identified as presenting satisfactory criteria (PBS scores from A to D), to be putatively involved in protein-protein interactions with MtNFP-kinase domain. The gene MtrunA17Chr3g0116591 scored D and was identified as belonging to the protein kinase RLK-Pelle-LRR-XI-2 family. Three clones containing partial sequences of this gene were detected in the Hybrigenics screening as interactants of NFP kinase domain (aa 319-429).

### Seed germination and growth conditions

Medicago seeds were scarified, sterilised and germinated as previously described (Luu et al. 2022).

### cDNA cloning

The full *SOBIR1* coding sequence (CDS) was amplified with Phusion High Fidelity DNA Polymerase (F530S, Thermo Fisher Scientific, USA) using *M. truncatula* A17 root cDNA as template and NEP013 primers (Table S1). The PCR product was cloned into pJet1.2/blunt using CloneJET PCR Cloning Kit (K1231, Thermo Fisher Scientific, USA). The correct clone was selected by sequencing, then used for further cloning steps.

### Constructs for in planta protein expression

For *Nicotiana benthamiana* leaf agro-infiltration, full length CDSs of SOBIR1 and NFP were fused at the C-terminus with the sequences encoding TagGFP and mCherry proteins, under the control of promoter Ubiquitin from *Lotus japonicus* (ProLjUbi) and the *Cauliflower mosaic virus* (CaMV) 35S promoter (Pro35S), respectively. The vector pCambia2200ΔDsRED was used (Fliegmann *et al*., 2016). In this experiment, the fusion of LYR3, a closely related protein of NFP, with mCherry was used as control (Fliegmann *et al*., 2016).

For *M. truncatula* root transformation, the NFP-ECD-TM were fused with three different phospho-versions of the NFP-KD (wild-type, phospho-silent and phospho-mimic) and were tagged at the C-terminus with 3xFLAG under the control of ProLjUbi by Golden gate cloning, using a vector based on pCAMBIA2200 expressing DsRED (Fliegmann *et al*., 2016). The empty vector (EV) was used as negative control.

### Transient expression of receptor constructs and the luciferase-reporter assay in Arabidopsis *protoplasts*

Transient co-transfection of the pFRK1:Luciferase reporter (Yoo, Cho, and Sheen 2007) with the receptor expression constructs in mesophyll protoplasts from *A. thaliana sobir1-12* mutant plants was performed as described (Wang, Albert, et al. 2016). Protoplasts were re-suspended in W5-solution containing 200 µM firefly luciferin (Synchem UG) and distributed in 96-well plates (20 000 protoplasts/90 µL). After overnight incubation at room temperature protoplasts were treated with nlp20 (10 times concentrated in W5-solution) or W5 as mock control and luciferase activity was recorded once per hour with a luminescence plate reader (Mithras LB 940, Berthold).

### Transient expression of receptor constructs and bioassays in *N. benthamiana*

Transient expression in *N. benthamiana* was performed as described (Albert et al. 2010). The oxidative burst was measured with leaf pieces floating on 100 µl water containing 20 µM L-012 (Wako) and 2 µg/ml horseradish peroxidase (Applichem), after addition of peptides, with a luminescence plate reader (Mithras LB 940, Berthold). The amount of ethylene was measured by GC in the headspace of 4 leaf pieces floating on 500 µl water, treated for 4 h with the peptides or controls (mock: 10 mg/ml BSA and 0,1 M NaCl). Expression of receptor constructs was verified by Western Blot analysis using anti-GFP, anti-Myc, or anti-HA antibodies and secondary antibodies coupled to alkaline phosphatase with CDP-Star (Roche) as substrate.

### Mutants and nodulation tests

*Tnt1* insertional mutants, *sobir1-1* (NF16427, containing 23 flanking sequence tags (FSTs)) and *sobir1-3* (NF20347, containing 35 FSTs) of *M. truncatula* R108, were obtained from the Noble Research Institute collection (USA) (now at Oklahoma State University, https://medicago-mutant.dasnr.okstate.edu/mutant/database.php), and *Mtsobir1-1* was backcrossed twice. In addition, *nad1-2* (NF14356) (Wang *et al*., 2016) was used for nodulation tests, and the double mutant *sobir1-1/nad1-2* was generated by crossing the *sobir1-1* mutant selected from the first backcross and the *nad1-2* mutant (Wang *et al*., 2016). All mutant lines were analysed in comparison to WT-like sister lines or R108 WT background.

For CRISPR/Cas9 gene editing, three guide RNAs were designed from the CDS of *MtSOBIR1* using CRISPOR (http://crispor.tefor.net/). The CRISPR modules were designed and combined as described (Jardinaud et al 2022). Transgenic *M. truncatula* 2HA plants were generated via *Agrobacterium tumefaciens, and r*egenerated plants were genotyped by PCR and sequencing using SOBcrp primers (Table S1). A 5-bp deletion mutation leading to an early stop codon forming a truncated protein (294 aa) was identified. After crossing out the Cas9, the mutant, *Mtsobir1-CRP*, was used for phenotyping. A sister line from the same transformation with no detected modification in *MtSOBIR1*, without Cas9 was used as control, or 2HA.

For *in vitro* tests, germinated seedlings were transferred to tubes or Petri dishes containing gelified Fahreus medium, 0.2 mM NH_4_NO_3_ (Luu et al., 2022). Alternatively, germinated seedlings were transferred into pots (8×8×8 cm), filled with sterilized attapulgite clay granules (Oil Dri, UK). To each pot, 80 ml of Fahraeus medium supplemented with 0.2 mM NH_4_NO_3,_ was added. After 3d (pots) or 6d (*in vitro*), plants were inoculated with approximately 10,000 bacteria/plant of *S. meliloti* 2011 WT or *S. medicae* WSM419. Plants were kept in growth rooms at 21/25°C, 16h photoperiod. Nodules were counted at different time points or at 21 dpi. Bacterial infection was determined by LacZ straining with X-gal (Arrighi et al 2006).

### Inoculation of Aphanomyces euteiches and assessment of the resistance level of the *M. truncatula lines*

Zoospores of the ATCC201684 *A. euteiches* strain, isolated from pea, were produced as previously described (Badreddine et al. 2008). Following germination, seeds of WT or mutant lines were grown *in vitro* on M medium (Becard and Fortin 1988) with a 16-h light at 22°C and 8-h dark at 20°C. Seedlings were inoculated one day after with a 5 µl droplet of zoospores (10^5^ sp./mL) deposited in the middle of the piliferous area of the roots. To assess resistance, three independent repeats containing at least 20 inoculated and 15 non-inoculated plants each, were performed. Percentages of root browning were measured and calculated at 14 and 21 days post inoculation using ImageJ software. The relative amount of *A. euteiches* was assessed by qRT-PCR (Gibelin-Viala et al. 2019).

### qRT-PCR

RNA samples were extracted using the Nucleospin plant RNA extraction kit (Macherey-Nagel GmbH & Co. KG, Germany). cDNAs were synthesized using PrimeScript RT Mastermix (Takara Bio Inc., Japan) and used as templates for qRT-PCR analysis using LightCycler 480 (Roche, Switzerland). Primers used for gene expression are listed in Table **S1**.

### Acetylene reduction assays

Assays were performed on inoculated plants at 21 dpi as described (Luu et al 2022). Briefly, 1 ml of acetylene was injected into a test tube containing one single inoculated plant and closed with a septum. Tubes were incubated in a growth chamber for two hours. Then, 400 µl of gas samples were analysed using a gas chromatograph equipped with a flame ionization detector (GC7820A, Agilent). Activity was normalized with the number of nodules per plant.

### Agro-infiltration and Fluorescence Lifetime Imaging on Nicotiana benthamiana leaves

*Agrobacterium tumefaciens LBA4404* strains containing ProLjUbi:SOBIR1-GFP or Pro35S:NFP-mCherry and LYR3-mCherry fusion constructs were used to agro-infiltrate the three to four oldest leaves of each *N. benthamiana* plant. At 3 dpi, the protein localisation was observed using confocal microscopy, and then sent to the FRAIB Imaging platform to perform FRET-FLIM measurements (described by Fliegmann et al., 2016).

### Kinase assays and identification of phosphosites

6xHistidine-Glutathione-S-transferase (GST) tagged proteins of the predicted intracellular region of SOBIR1 (termed the KD) was cloned into pCDF-Duet vector (Novagen, EMD Chemicals, San Diego, CA) and expressed in *E. coli* Rosetta/DE3 and the proteins purified using glutathione resin (GE Healthcare, USA) as described (Fliegmann *et al*., 2016). SOBIR1-KD was released from the resin using PreScission Protease (GE27-0843-01, Sigma Aldrich, Germany). SOBIR1-KD was incubated with kinase buffer containing [γ-^32^P]-ATP either alone or with purified GST/NFP-KD, GST/LYR3-KD, GST/LYR4-KD, GST/LYK3-deadKD (G334E mutation), Myelin Basic Protein (MyBP) or GST at 25⁰C for 1 h and the proteins analysed by SDS-PAGE, followed by Coomassie staining and Phosphor Imaging.

The identification of phosphosites was done using Phos-tag gel electrophoresis to separate phosphorylated and unphosphorylated proteins followed by excision of the appropriate bands and in-gel protease digestion. Peptides were prepared and analysed by mass spectrometry (MS) by the GenoToul Proteomic platform as described by Fliegmann et al. (2016).

### Medicago complementation assays

Using *Agrobacterium rhizogenes*-mediated transformation (Boisson-Dernier et al. 2001) seedlings of *nfp-2* (Arrighi *et al*., 2006) and wild-type A17 were transformed using strains containing either empty vector (EV) or different versions of ProLjUb:NFP-ECD-TM/NFP-KD-3xFLAG constructs, and transformed roots were selected on medium containing 25 µg.ml^-1^ kanamycin, and after two-weeks growth, by expression of the DsRed marker. Nodulation was analysed in pots as above after 4 wk. The number of nodulated plants and the number of nodules/nodulated plant were analysed.

### MtSOBIR1 *p*romoter study

A 3kb genomic region upstream of the ATG of *MtSOBIR1* was cloned from genomic DNA of *M. truncatula* A17 using the proSOBIR1_GG primers (Table S1). The promoter was cloned with the CDS of *GUS (β-glucuronidase)* using the Golden Gate cloning method and transformed into A. *rhizogenes* Arqua1 as described by Fliegmann et al. (2016). Hairy root transformation and GUS staining were performed as previously described (Boisson-Dernier *et al*., 2001, Arrighi et al 2006). Nodule sections were made before GUS staining.

## Results

### MtSOBIR1 is a putative interacting partner of MtNFP

MtSOBIR1 was found in a yeast two-hybrid screen using the MtNFP kinase domain (NFP-KD) as bait to a cDNA library of *M. truncatula* plants inoculated, or not, with the legume oomycete pathogen *Aphanomyces euteiches*. Interactions between the bait and prey clones were re-verified in “one by one” tests on solid growth selective media containing or lacking Histidine (Fig. S1). *In silico* analysis of the protein encoded by the *MtSOBIR1* gene (MtrunA17_Chr3g0116591) predicts an LRR-RLK that is highly similar to AtSOBIR1 (61% identity and 75% similarity; Fig. S2). Both proteins contain five LRRs in the ECD, an extracellular juxtamembrane (eJM) domain, a GxxxGxxxG motif in the TM that is implicated in protein interactions (Bi *et al*., 2015), and a KD with conserved features of active kinases; a P loop, an activation loop and HRD and DFG motifs. Phylogeny analysis indicated that MtSOBIR1 forms a clade with SOBIR1 homologs in other plants, but not with the the CRN sub-clade of LRR-RLK XI-2 proteins, and not with the next closest *M. truncatula* protein (MtrunA17_Chr1g0183231; Fig. S3). Except for the allotetraploid species *Glycine max, M. truncatula* like most plant species contains only one copy of SOBIR1 in this analysis. Together, this analysis indicates that MtSOBIR1 is the unique ortholog of AtSOBIR1. It is because of these results and subsequent functional analysis, that we have adopted the name MtSOBIR1.

### MtSOBIR1 partially complements *Arabidopsis thaliana sobir1*

To determine whether MtSOBIR1 is functionally equivalent to AtSOBIR1, we tested whether MtSOBIR1 could replace AtSOBIR1 for the formation of an active complex with AtRLP23. For this, mesophyll protoplasts from *Atsobir1-12* were co-transformed with the reporter construct pFRK1::luciferase, and the expression construct for the AtRLP23 receptor, AtRLP23-GFP, and, in addition, an expression construct, either for MtSOBIR1-GFP or for AtSOBIR1-Myc. These cells were treated with 1 µM of the ligand of AtRLP23, nlp20, and luciferase activity was measured for 6 hours. A clear increase in activity was detected for AtSOBIR1-expressing cells and, although to a lesser degree, also for MtSOBIR1-expressing cells, but not in protoplasts that only expressed AtRLP23 (Fig. 1). We also transiently expressed MtSOBIR1 in a CRISPR/CAS *Nicotiana benthamiana* mutant together with AtRLP23, and observed increases in ethylene and ROS production upon treatment with nlp20, similarly to AtSOBIR1 (Fig. S4). Together, this indicates that MtSOBIR1 can functionally replace AtSOBIR1 as a positive regulator of immunity.

**Figure 1.**
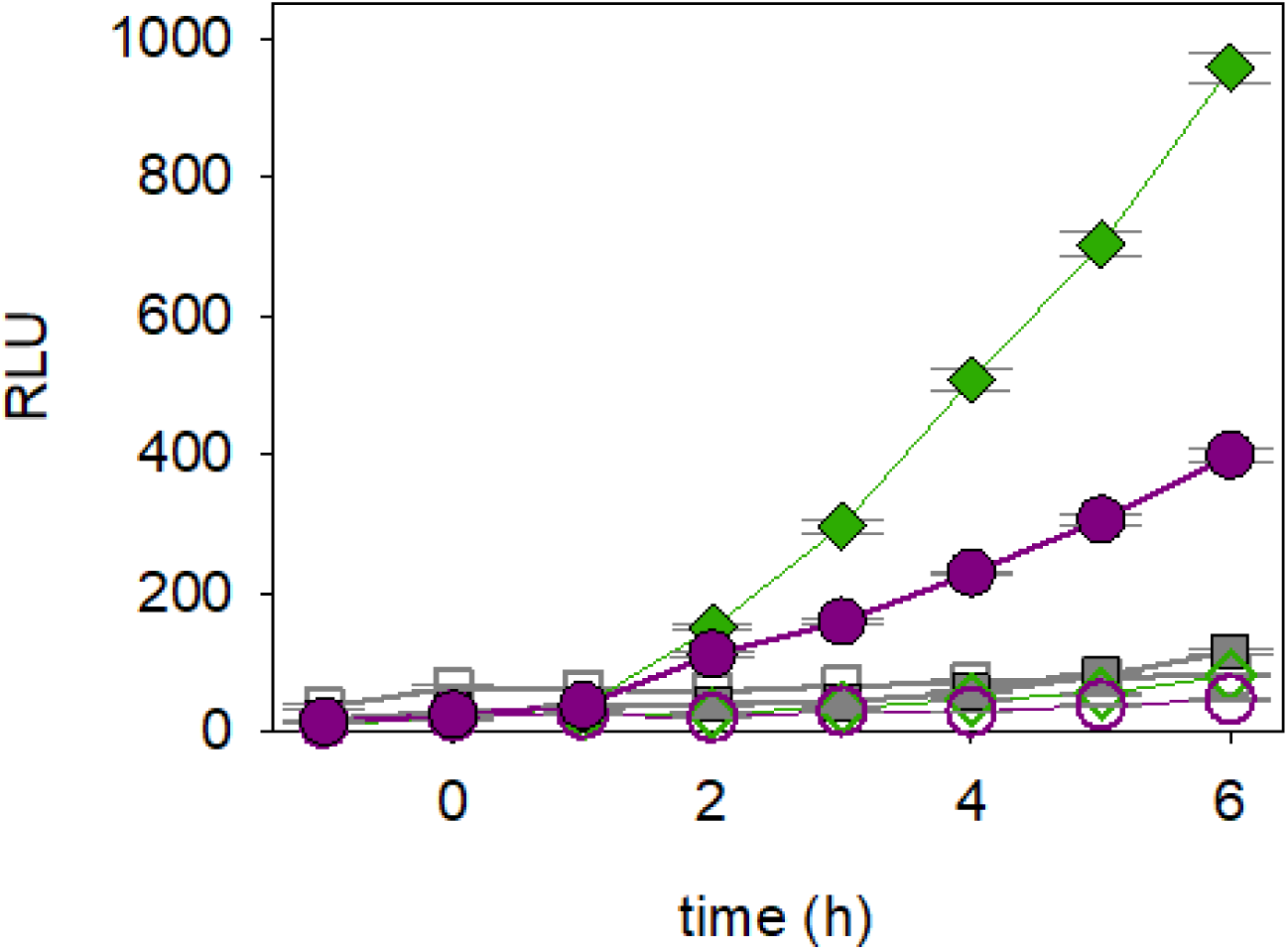
Complementation of *A. thaliana sobir1-12* by MtSOBIR1. Mesophyll protoplasts from *A. thaliana* lacking SOBIR1 (*sobir1-12*) were co-transformed with pFRK1::luciferase and the expression construct for AtRLP23-GFP (gray symbols), and in addition either the expression construct for MtSOBIR1-GFP (pink symbols) or the one for AtSOBIR1-Myc (green symbols). Treatment with 1 µM nlp20 (filled symbols) at the time point 0 (arrow) resulted in an increase of luciferase activity (RLU, relative light units) in Mt- or AtSOBIR1-expressing cells, but not in protoplasts which expressed RLP23 only. Values and error bars indicate mean and standard deviation of 4 replicates.

### Validation of the physical interaction between MtSOBIR1 and MtNFP

To validate the results of the Y2H screen, the MtNFP-MtSOBIR1 interaction was analysed by FRET-FLIM. The full-length CDS of MtSOBIR1 in *M. truncatula* A17 was fused with TagGFP, under the control of the Ubiquitin promoter from *L. japonicus* (pLjUbi). The protein was transiently expressed individually or co-expressed with an NFP-mCherry fusion in *N. benthamiana* leaves by *Agrobacterium tumefaciens*-mediated transformation. The co-expression of MtSOBIR1-GFP with MtLYR3-mCherry, a closely-related protein to MtNFP, was used as a control (Fliegmann *et al*., 2016).

At 3 dpi, confocal microscopy showed that MtSOBIR1 is detected at the periphery of *N. benthamiana* leaf cells and co-localises with either MtNFP or MtLYR3 (Fig. 2a, b). As MtNFP and MtLYR3 were previously reported to localise at the plasma membrane of *N. benthamiana* leaves (Lefebvre *et al*., 2012; Fliegmann *et al*., 2016), we conclude that the peripheral cellular location of MtSOBIR1 is at the plasma membrane.

**Figure 2.**
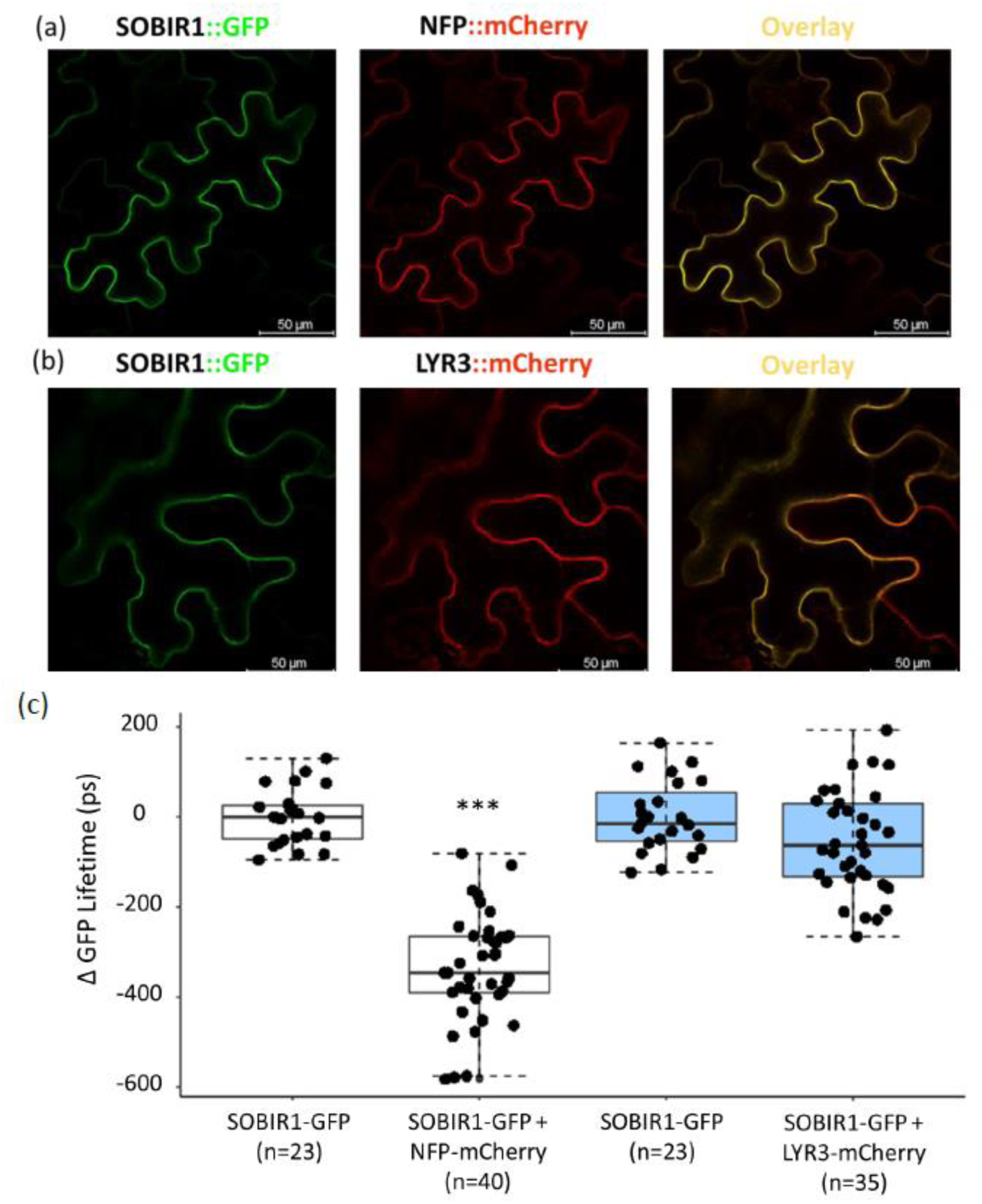
Validation of the physical interaction between MtSOBIR1 and MtNFP. Co-localisation of MtSOBIR1 and MtNFP (a) and MtSOBIR1 and MtLYR3 (b) at the plasma membrane of *N. benthamiana* leaves. Leaf discs were observed by confocal microscopy at 3 dpi. Bars, 50μM. (c) FRET-FLIM experiments between MtSOBIR1 and MtNFP, MtLYR3 at the plasma membrane of *N. benthamiana* leaves. For each measurement, the difference to average GFP decay times (Δ) of the SOBIR1-GFP in the absence of the mCherry fusions is given in ps; n: number of samples measured for each condition. Different colours indicate separate experiments. Students’ *t*-tests were used for statistical analyses (***, *P* < 0.001).

Similar leaf samples were used for FRET FLIM analysis in which MtSOBIR1-GFP was used as the donor and either NFP-mCherry or MtLYR3-mCherry was the acceptor. The co-expression of MtSOBIR1and MtNFP led to a significant reduction of GFP lifetime compared to MtSOBIR1 alone, whereas no such change was observed when MtSOBIR1 was co-expressed with MtLYR3 (Fig. 2c). This indicates a physical interaction between MtSOBIR1 and MtNFP, which is specific in comparison with the closely related LysM-RLK MtLYR3.

### MtSOBIR-KD autophosphorylates and trans-phosphorylates MtNFP-KD on multiple residues

To test whether MtSOBIR1 has an active KD and can trans-phosphorylate the pseudo-kinase domain of MtNFP, *in vitro* phosphorylation assays on the intracellular regions of the two proteins (which includes the KDs), purified after expression in *E. coli* were performed. The ability of MtSOBIR1 to trans-phosphorylate related proteins, MtLYR3-KD, MtLYR4-KD and MtLYK3-deadKD (Fliegmann et al. 2016), and the model kinase substrate myelin basic protein (MyBP) was also tested.

MtSOBIR1-KD had an autophosphorylation activity and could trans-phosphorylate a GST fusion of MtNFP-KD and also MyBP in the presence of [γ-^32^P] ATP, but not GST and BSA (Fig. 3). In addition, MtSOBIR1-KD was able to trans-phosphorylate other related pseudo kinases including MtLYR3-KD and MtLYR4-KD, and MtLYK3-deadKD (Fig. S5). Phosphorylated and unphosphorylated MtSOBIR1-KD were separated on a Phostag gel (Fig. S6). The phosphorylated proteins were then extracted, in-gel digested with Trypsin, and analysed by nano-liquid chromatography–tandem mass spectrometry (LC–MS/MS). Two different samples including JC01 (SOBIR1-KD alone) and the JC02 (SOBIR1-KD in the presence of MtNFP-KD) were analysed. Twenty-five phosphosites including both Ser/Thr and Tyr were identified (Table S2). Only seven out of 25 sites are conserved with the 14 sites identified in AtSOBIR1-KD (Mitra *et al*., 2015) (Fig. S7). In addition, there are 8 sites that were detected in JC01 but were not found in JC02 suggesting a possible inhibitory role of MtNFP-KD on the autophosphorylation activity of MtSOBIR1-KD.

**Figure 3.**
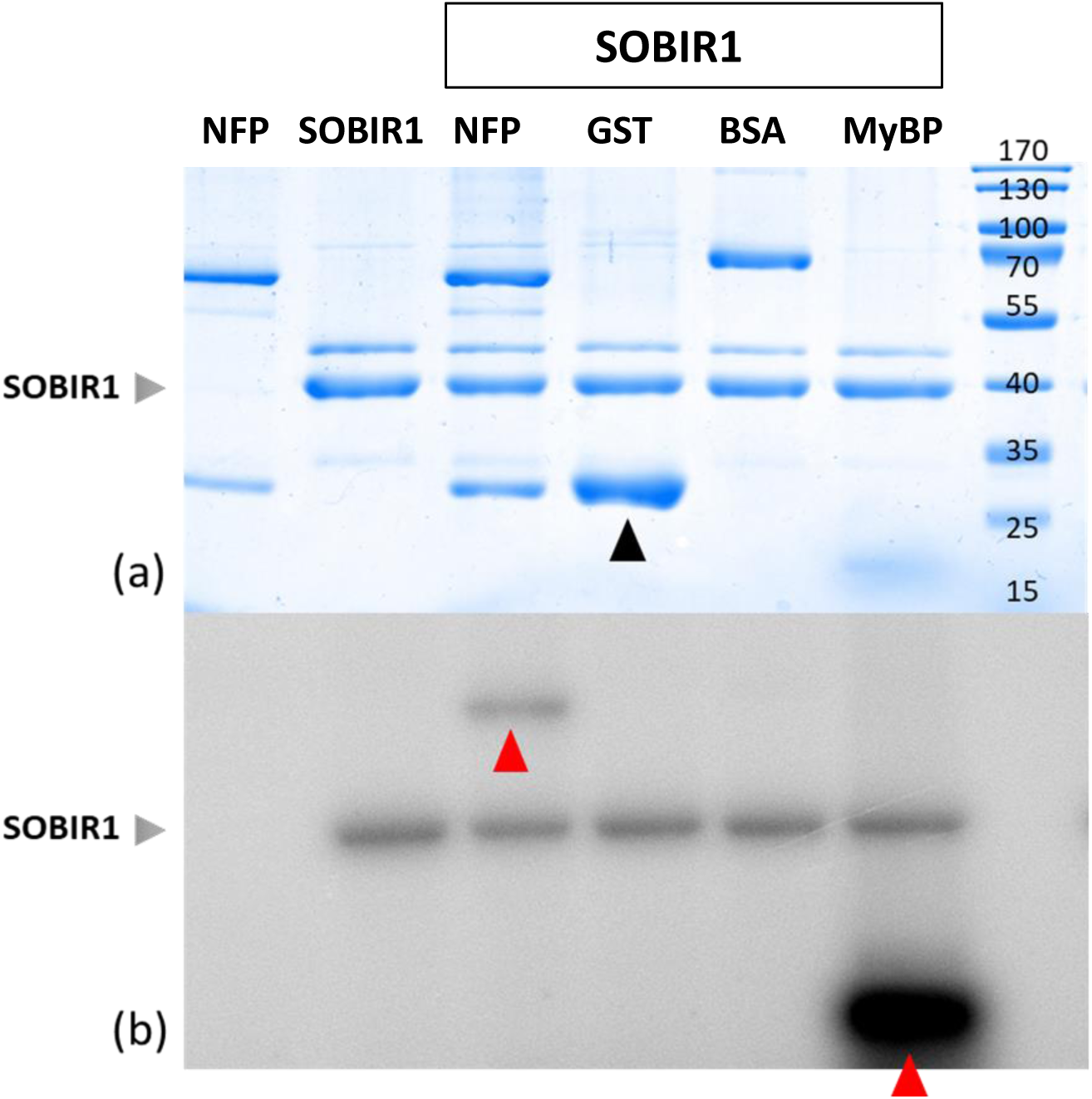
*In vitro* autophosphorylation and transphosphorylation activity of the kinase domain of MtSOBIR1 (MtSOBIR1-KD). The KDs of MtNFP (lane 1) and MtSOBIR1 (lane 2) were expressed and purified from *E. coli* as fusions wth 6His-GST and GST, respectively. Soluble 6His-GST-cleaved SOBIR1-KD was incubated alone (lane 2) or with purified GST-NFP-KD, GST, BSA and MyBP (lanes 3-6) in the presence of radioactive [ɣ-32P] ATP at 25ᵒC for 1h. Assays were analysed by SDS-PAGE, followed by a Coomassie staining (a) and phosphor imaging (b). Transphosphorylated proteins are marked by red arrowheads on the phosphor-image. The position of GST is marked on the Coomassie gel (black arrowhead).

A similar approach was used to identify the phosphosites in MtNFP-KD that were trans-phosphorylated by MtSOBIR1-KD (Fig. S6). Four different samples were used for the analyses. In JC03 where there is only GST and MtNFP-KD, no phosphosites were identified. In other samples, there are differences in the number of phosphosites found in each sample, specifically, six sites in JC04, nine sites in JC05 and four sites in JC06 (Table S3). In total, nine sites were identified with the frequency of detection within the 3 samples indicated (Table S3). By comparison to the ortholog of MtNFP in *L. japonicus*, LjNFR5, which is trans-phosphorylated by both LjNFR1-KD (1 site) and LjSYMRK-KD (8 sites) (Madsen *et al*., 2011), only 1 phosphorylated site of MtNFP-KD is conserved, corresponding to a site of LjNFR5-KD trans-phosphorylated by LjSYMRK-KD (Fig. S8).

To investigate the possible biological function of the MtNFP-trans-phosphorylated sites, two phospho-versions of MtNFP-KD were generated: the phospho-silent version (all 9 sites were mutated to Ala – 9A) and the phospho-mimic version (all 9 sites were mutated to Asp – 9D) (Fig. S9). These KDs, as well as the WT MtNFP-KD, were fused with the WT NFP-ECD-TM domains and tagged with 3xFlag under the regulation of ProLjUbi, and then transformed via *A. rhizogenes* into *Mtnfp-2* plants. The *Mtnfp-2* mutant is strictly Nod minus (Arrighi et al 2006). Following inoculation with *Sinorhizobium meliloti* 2011, all the NFP constructs could restore nodulation to the *Mtnfp-2* mutant to a similar extent, while the EV did not (Fig. S9). This indicates that the phosphosites, identified by MtSOBIR1 transphosphorylation, are not essential for MtNFP function in nodulation, although we cannot exclude that they play some role in different conditions.

### Does MtSOBIR1 play a role in symbiosis and/or pathogen immunity?

*In silico* analyses of *M. truncatula* transcriptomic data in MtExpress (Carrere, Verdier, and Gamas 2021), showed that *MtSOBIR1* is expressed throughout plants (leaves, shoots and roots), including good expression in the root epidermis and nodules (Fig. S10). By RT-qPCR we showed that *MtSOBIR1* expression in roots is regulated in response to the oomycete pathogen *Aphanomyces euteiches* (Fig. S11). By transforming a transcriptional promoter-*MtSOBIR1*-GUS fusion into *M. truncatula* WT roots, we observed promoter activity in roots, including root hairs, and in nodules formed by *S. meliloti* 2011 (Fig 4). Nodule sections revealed a pattern in the apices of young nodules, as well as in vascular tissue and tissue around the nodule vasculature (Fig 4). Promoter-*MtSOBIR1*-GUS expression decreased in older nodules, apparently disappearing from nodule apices (Fig 4).

**Figure 4.**
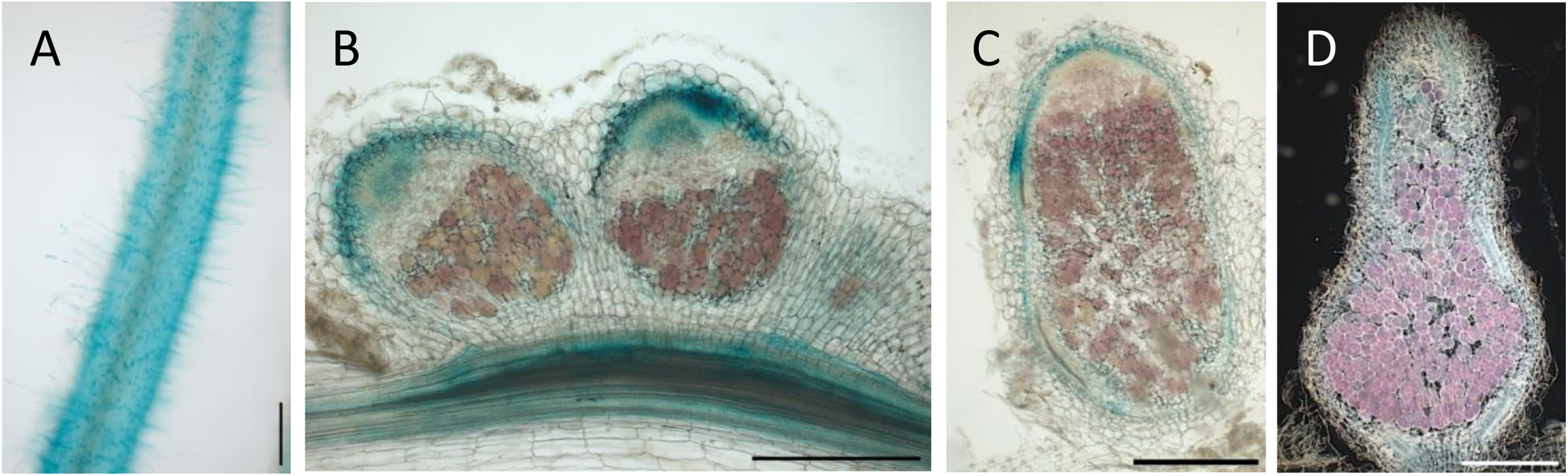
Spatio-temporal expression analysis of pMtSOBIR1-GUS in roots and nodules following inoculation with *S. meliloti* 2011. A, roots 10 dpi showing expression in root hairs B-D, sections of nodules at different stages of development. Approximate nodules ages: 29 days (B, C), 56 days (D). Scale bars, 250 µm

Genomic data analysis identified a single *SOBIR1* sequence in the contrasting *M. truncatula* genotype R108, as for A17. The A17 and R108 proteins share about 99% identity, suggesting no significant variation of this protein in the two genotypes. To study the role of *MtSOBIR1*, we identified two allelic *Tnt1* insertion mutants (*Mtsobir1-1* and *Mtsobir1-3*) in the R108 background, and we generated a CrispR-cas9 mutant in the *M. truncatula* 2HA background (Fig. S12). All 3 mutants are predicted to be knock-outs of SOBIR1 function, which we verified for *Mtsobir1-1* and *Mtsobir1-3* (Fig. S12). The WT sibling lines were selected as controls.

#### A role for *MtSOBIR1* in immunity against *Aphanomyces euteiches*?

Given that SOBIR1 orthologs can play positive roles in immunity against pathogens, we tested whether this is also the case for MtSOBIR1. The cDNA library used to identify MtSOBIR1 was made from *M. truncatula* roots infected or not by *Aphanomyces euteiches.* This is a well-studied oomycete pathogen of *M. truncatula* (Djebali et al. 2009), and shows enhanced virulence on *Mtnfp* mutants (Rey et al 2013). We therefore studied the ability of *Mtsobir1* mutants to respond to and to interact with *A. euteiches*.

Firstly, we measured defence gene expression of *Mtsobir1* roots in response to *A. euteiches*. We took *Mtsobir1-1* mutant and WT root samples 24 h after inoculation by *A. euteiches* and performed RT-qPCR analysis on 5 immunity-associated genes known to be rapidly induced in *M. truncatula* by *A. euteiches* (Rey et al 2013) (Rey et al. 2016) (Rey et al. 2019). In WT *MtSOBIR1-1* roots we confirmed the rapid induction of four of these genes, encoding a CHalcone Synthase (MtCHS, MtrunA17_Chr7g0220181), a Pathogenesis - Related protein (MtPR10; MtrunA17_Chr4g0067951), a peroxidase (MtPRX67, MtrunA17_Chr4g0043391) and a Respiratory Burst Oxidase Homolog (MtRBOHD; MtrunA17_Chr3g0131361) (Fig. 5A). Of these, the MtPR10, MtPRX67 and MtRBOHD genes were not induced in mutant *Mtsobir1-1* roots (Fig. 5A), indicating a role for MtSOBIR1 in the ability of *M. truncatula* to respond to *A. euteiches* by activating a defence response. The other 2 genes, the MtCHS and the thaumatin (MtrunA17_Chr1g0180221), did not follow this pattern (Fig 5A). Finally, the symbiotic gene *MtSymCRK*, which has a role in repressing defence responses in nodules (Berrabah et al 2014), was not induced in WT roots, as we expected, but could have a higher level in mutant *Mtsobir1-1* roots compared to WT (Fig. 5A).

**Figure 5.**
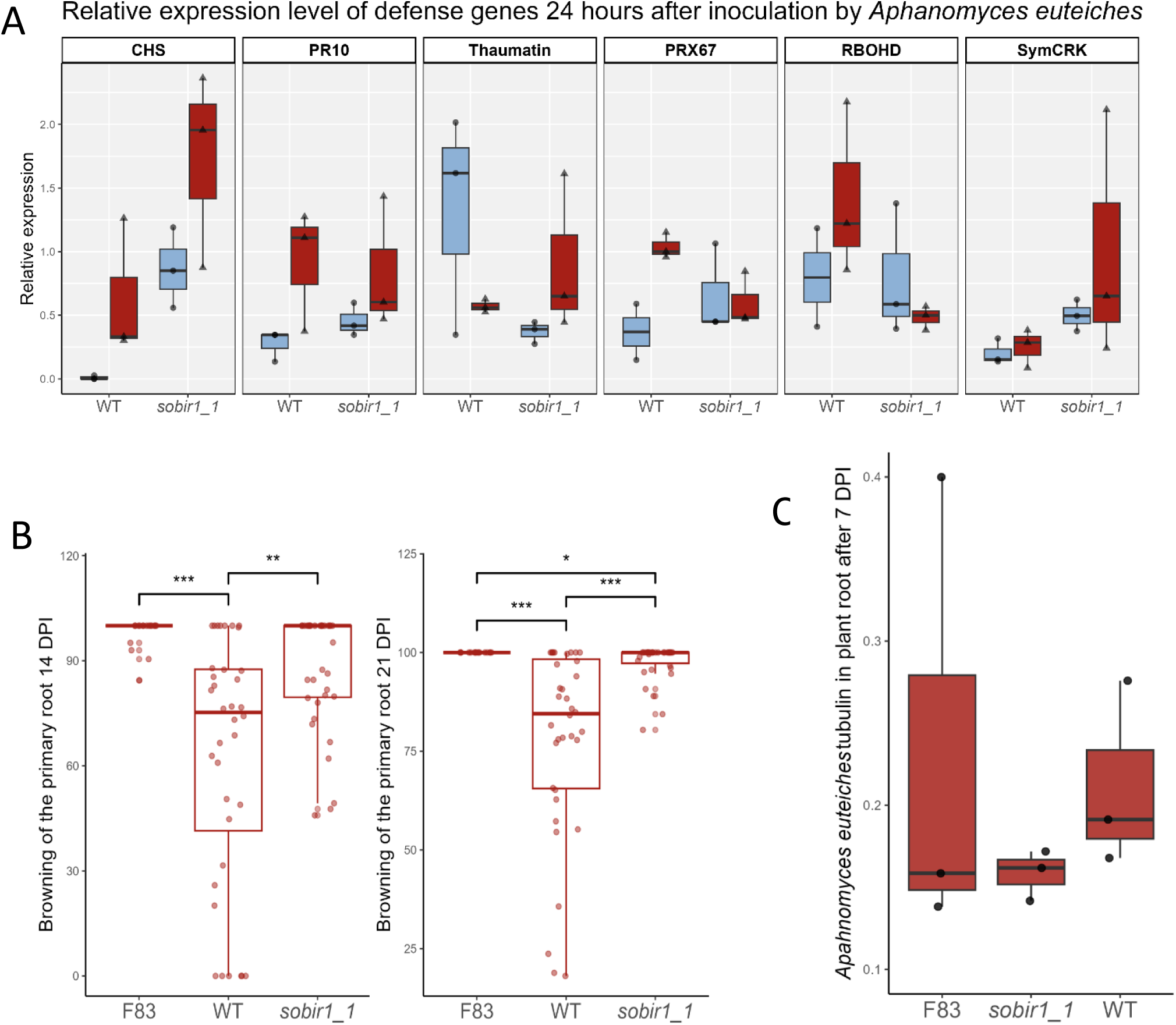
*Mtsobir1-1* mutant phenotypes in response to *Aphanomyces euteiches.* A, Measurements by RT-qPCR of defence gene induction by *A. euteiches* in WT (*Mtsobir1-1* WT) and *MtSobir1-1* plants 24 hpi (blue = non-inoculated; red = inoculated). B, Symptom development at 14 and 21 dpi in *M. truncatula* F83 (susceptible genotype), WT (*Mtsobir1-1* WT) and *MtSobir1-1,* plants following *A. euteiches* inoculation. C, Measurements by RT-qPCR of *A. euteiches* tubulin gene expression in inoculated *M. truncatula* F83 (susceptible genotype), WT (*Mtsobir1-1* WT) and *MtSobir1-1* plants 7 dpi. * (P < 0.05), ** (P < 0.01), *** (P < 0.001)

We then performed disease susceptibility tests on *in vitro* grown seedlings, with symptom development assessed by measuring root browning and the susceptible *M. truncatula* genotype F83 used as a control (Djébali et al 2009). These tests indicated that the *Mtsobir1-1* mutant was more susceptible than its WT sibling line at 14 and 21 dpi (Fig. 5B), but when *A. euteiches* was quantified in infected *Mtsobir1-1* mutant roots, as another possible indicator of susceptibility, no difference was found to WT plants (Fig. 5C). The *Mtsobir1-3* mutant was also more susceptible than its WT control line, but no differences were observed between the *Mtsobir1-CRP* mutant and its WT line (Fig. S13). This indicates that *MtSOBIR1* plays a role against *A. euteiches* in the *M. truncatula* R108 genotype.

#### A symbiotic role for *MtSOBIR1*?

*Mtsobir1* mutants were tested for nodulation with *S. meliloti* 2011, a well-studied symbiont of *M. truncatula*. No differences were found between any of the mutants and their WT siblings in nodule numbers, nitrogenase activity or nodule structure (Fig. S14). Infection, nodulation kinetics and percentages of plants nodulated were not different either between mutant and WT lines (Fig. S15), indicating that *MtSOBIR1* is not important for interaction with *S. meliloti* 2011. Efficient symbiotic nitrogen fixation by rhizobia relies on the control of plant immunity, (Berrabah *et al*., 2023). The *Mtnad1-2* mutant shows brown, necrotic, non-nitrogen fixing (Fix^-^) nodules in which numerous defence genes are expressed, unlike in WT nodules (Wang *et al*., 2016). To find out whether *MtSOBIR1* might intervene in the modulation of defence responses inside nodules via the *NAD1* pathway, we generated a double *Mtsobir1-1/Mtnad1-2* mutant, and tested nodulation with *S. meliloti* 2011. The double *Mtsobir1-1/Mtnad1-2* mutant did not show any significant difference in either nodule numbers, nodule types or nitrogenase activity compared to the *Mtnad1-2* single mutant (Fig. S16). Besides, the WT sibling *MtSOBIR1-1/MtNAD1-2* showed a clear rescued phenotype in both categories compared to the *Mtnad1-2* mutant. This result suggests that *MtSOBIR1* does not intervene in the modulation of defence responses inside nodules via the *NAD1* pathway.

Since plant symbiotic phenotypes can be dependent on rhizobial strain, we tested *Mtsobir1* mutants with the *S. medicae* strain WSM419 that changes symbiotic phenotypes of some plant genotypes compared to *S. meliloti* 2011 (Jardinaud et al. 2023) (Luu et al 2022). Nodulation tests in the 2HA genotypes with *S. medicae* WSM419 showed no differences between the CrispR-cas9 *Mtsobir1* mutant and its WT control (Fig. 6A). However, both R108 mutants, *Mtsobir1-1* and *Mtsobir1-3,* showed poor nodulation with *S. medicae* WSM419 compared to their WT sibling lines (Fig. 6A, Fig. S17). As before, no differences were seen in nodule numbers between mutant and WT control plants with *S. meliloti* 2011, while *Mtsobir1-1* and *Mtsobir1-3* mutants nodulated better with *S. meliloti* 2011 than with *S. medicae* WSM419 (Fig. 6A).

**Figure 6.**
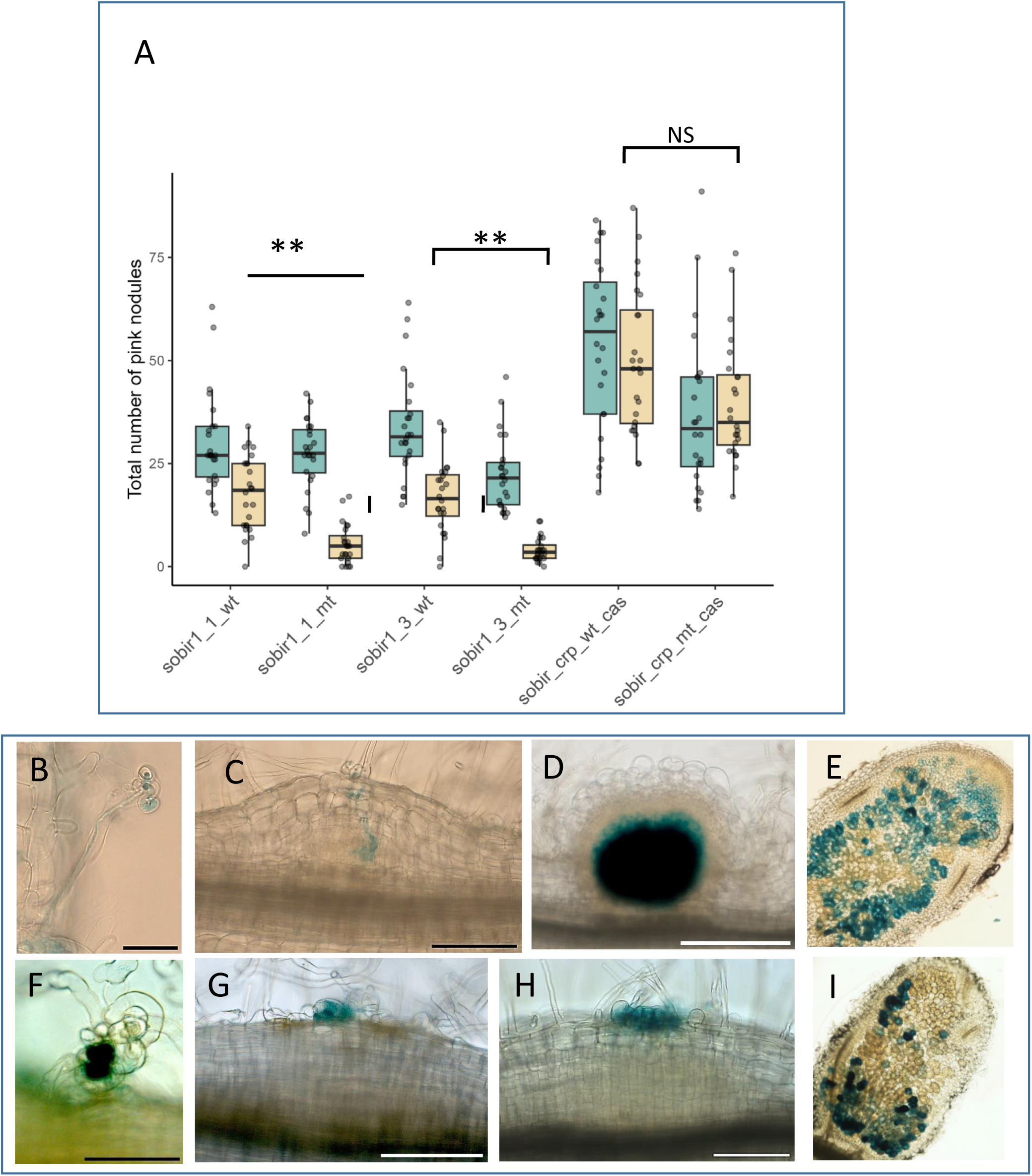
*Mtsobir1* mutants in the R108 background nodulate badly with *Sinorhizobium medicae* WSM419. A, Nodulation tests with mutant and WT genotypes as indicated, and with both *S.meliloti* 2011 (green) and *S. medicae* WSM419 (orange). Nodules were counted at 4 weeks PI in pot-grown plants. ** (P < 0.01), NS, not significant. B-I, Photos illustrating the infection phenotypes of WT and *Mtsobir1* mutants with *S. medicae* WSM419; B-E WT (R108); F-I *Mtsobir1* mutants. *S. medicae* WSM419 is coloured in blue by lacZ staining. B, 10dpi; C 10 dpi; D; 14 dpi; E, 3wpi; F, *Mtsobir1-3* 10dpi; G, *Mtsobir1-3* 14dpi H, *Mtsobir1-1* 21dpi; I *Mtsobir1-1* 21dpi. Scale bars 50 µm (B, F), 100 µm (C, D, G, H).

*Mtsobir1-1* and *Mtsobir1-3* mutants inoculated with *S. medicae* WSM419 showed areas of uninfected cortical cell division, including at the bases of lateral roots, in addition to the rare nodules (Fig. 6). Above the areas of uninfected cortical cell division, rhizobia were abnormally trapped in root hairs, with no infection threads (Fig. 6F, G, H). Sections of nodules formed on *Mtsobir1-1* plants by *S. medicae* WSM419, did not show obvious differences with WT nodules formed by *S. medicae* WSM419 (Fig.6E, I).

Since ethylene is a well-known inhibitor of nodulation and implicated in nodule defence mechanisms (Penmetsa and Cook 1997) (Berrabah et al. 2018), we tested nodulation of the *Mtsobir1-1* mutant and its WT sibling line with *S. medicae* WSM419 in the presence of the ethylene inhibitor aminoethoxyvinylglycine (AVG), at 100 nM. This showed that AVG accelerated nodulation of the WT line and improved nodulation of *Mtsobir1-1* plants, but did not restore nodulation to the WT level (Fig. S18).

Nodule activity measurements of *S. medicae* WSM419-induced *Mtsobir1* mutants showed that *Mtsobir1-1* mutant and WT plants did not differ in their nitrogenase activities (Fig. S19). No difference was detected either between *S. medicae* WSM419 and *S. meliloti* 2011-induced *Mtsobir1* mutant nodules (Fig. S19). Despite these similarities, we compared gene expression in nodules formed by *S. medicae* WSM419 and *S. meliloti* 2011. To avoid contamination and to have the same conditions used for the nitrogenase activity tests, we produced nodules *in vitro*. The relative poor efficiency of *S. medicae* WSM419 on R108 compared to A17 is associated with early senescence of nodules (Kazmierczak et al. 2017). To study whether these features were worse in *S. medicae* WSM419-induced nodules formed on R108 *Mtsobir1* plants, we measured by RT-qPCR the expression of 3 cysteine protease genes, markers for nodule senescence, CP1, CP5 and CP6 (Sauviac et al. 2022). Of these, CP1 and CP6 were more highly expressed in *Mtsobir1-1* - *S. medicae* WSM419 nodules compared to WT and compared to *Mtsobir1-1* – *S. meliloti* 2011, indicating a higher level of senescence (Fig 7). Because of the importance of the suppression of plant defence gene expression for successful bacteroid persistence (Berrabah et al. 2023), although we had found no evidence for a role of MtSOBIR1 in the *NAD1* pathway for defence suppression, we also measured the expression of marker genes for nodule defence. We chose the same genes tested in *Mtsobir1* mutant plants in response to *A. euteiches* (Fig. 5), as well as genes whose repression in nodules is controlled by the nodule immunity genes *MtDNF2*, MtSYMCRK and *MtNAD1* (Berrabah et al 2023). RT-qPCR analysis showed higher levels for chalcone synthase, PR10 and NDR1 (Non-race specific Disease Resistance 1) in *Mtsobir1-1* - *S. medicae* WSM419 nodules compared to both WT-*S. medicae* WSM419 nodules and *Mtsobir1-1* – S. meliloti 2011 nodules (Fig. 7). This indicates a role of MtSOBIR1 in suppressing defence responses in nodules. The expression of the nodule immunity gene *MtSYMCRK*, was not modified in mutant nodules (Fig. 7).

**Figure 7.**
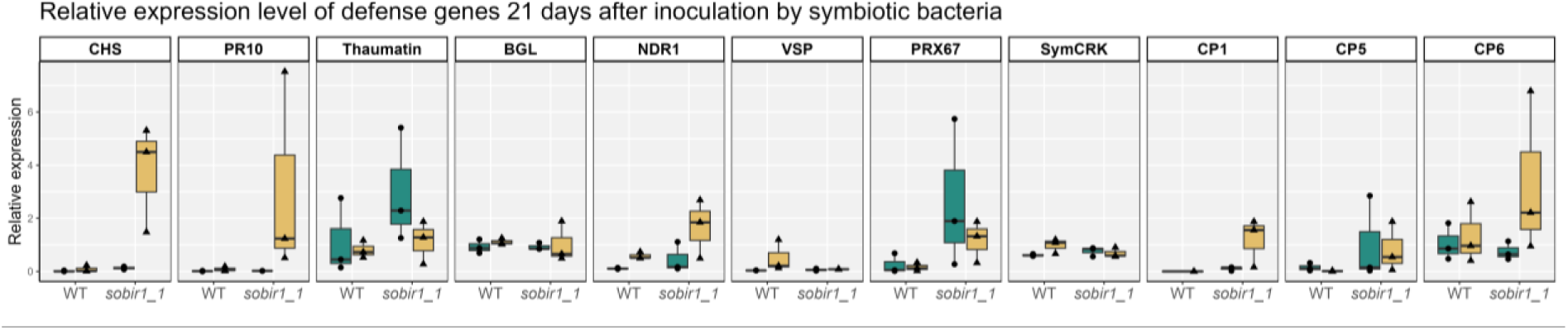
*MtSOBIR1* negatively regulates defence and senescence gene expression in *S. medicae* WSM419 nodules. Q PCR analysis of expression of marker genes for defence and nodule senescence in 21 d old nodules of *Mtsobir1-1* and R108 (WT) nodules formed by either *S. meliloti* 2011 (green) or *S. medicae* WSM419 (orange).

Finally, another phenotype was observed for *Mtsobir1-1* and *Mtsobir1-3* mutants when grown and inoculated in *in vitro* conditions. In response to both *S. meliloti* 2011 and *S. medicae* WSM419, many epidermal root cells of *Mtsobir1-1* and *Mtsobir1-3* plants appeared brown and necrotic (Fig. S20). In some cases, these cells were adjacent to aborted infection events (Fig. S20). This re.sponse suggested the presence of Reactive Oxygen Species (ROS) and oxidised phenolics in these cells, as in a Hypersensitive Reaction (HR). However, these cells did not colour with Evans Blue, a marker used to detect HR in leaves, possibly due to differences between roots and leaves, for example thickened cell walls around such cell death in roots.

## Discussion

Establishment of a successful rhizobial legume symbiosis depends on activation of symbiotic signaling pathways in the host plant. These recruit components of endogenous developmental and hormonal pathways, and success depends on tight controls of specificity and defence reactions. There are parallels between symbiont and pathogen-triggered responses, including changes in levels of ROS, which, when altered, can lead to early blocks in rhizobial infection (Montiel et al. 2012). Evidence for overlaps in symbiont and pathogen-triggered responses, is also provided by the symbiotic plant mutants that are affected in their interactions with pathogens (Rey and Jacquet 2018). *MtNFP* is a good example of a plant gene with a dual role in symbiosis and plant immunity (Gough and Jacquet 2013). However, although the symbiotic roles of MtNFP have been characterised (including Nod factor perception, rhizobial infection and nodule organogenesis) (Arrighi et al 2006), less is known about how MtNFP controls plant immunity. In other plant species, notably *A. thaliana*, a major class of plant immunity receptors are LRR-proteins, both with (LRR-RLKs) and without (LRR-RLPs) a kinase domain, respectively. SOBIR1 is a good example of an LRR-RLK controlling immunity against plant pathogens. Knock out or knock down *sobir1* plants are often more susceptible to oomycete and fungal pathogens, such that the incompatibility of these interactions is dependent on SOBIR1. In this work, we have characterised MtSOBIR1 as an interactor of MtNFP, and as a new symbiotic component that is apparently needed for full compatibility of symbiont recognition in the *M. truncatula* R108 – *S. medicae* WSM interaction.

### The MtSOBIR1-MtNFP interaction

We showed that MtSOBIR1 co-localises and physically interacts with MtNFP at the plasma membrane of *N. benthamiana* leaves (Fig. 2). We also showed that MtSOBIR1-KD can trans-phosphorylate the KD of MtNFP on 9 sites (Table S3), although complementation analysis of phospho-dead and phospho-mimic versions of MtNFP for these 9 sites indicated that they are dispensable for nodulation (Fig. S9). Of these 9 sites, one is trans-phosphorylated on LjNF5-KD (the *L. japonicus* ortholog of MtNFP) by the symbiotic LRR-RLK LjSYMRK (Madsen et al. 2011). Some of the other phospho-sites on LjNFR5-KD identified with LjSYMRK-KD, as well as 9 predicted phospho sites on MtNFP, have all been shown to be nonessential when tested in complementation tests (Madsen et al. 2011) (Lefebvre et al. 2012), underlining the difficulty of finding biologically relevant phospho sites in these proteins using these techniques.

SOBIR1 proteins are well known co-receptors, and their roles generally involve complex formation with other receptors, notably RLPs (Gust and Felix 2014) (Snoeck et al 2023), although some SOBIR1-RLCK interactions have also been reported (Pruitt et al. 2021). SOBIR1 interactors are generally involved in immunity, but SOBIR1 can also interact with development-related RLPs such as CLV2 and TMM (Liebrand et al 2014), and, although the mechanism is not clear, SOBIR1 is a negative regulator of the HAES/HAESA-like2 pathway for organ abscission (Gubert and Liljegren 2014). Gx_3_G motifs in the TM of SOBIR1 proteins, and in interacting RLPs, are important for the interactions of these proteins (Bi et al. 2016); (Snoeck et al 2023). This region of MtSOBIR1 is highly conserved with that of AtSOBIR1 (Fig. S2), and partially with the TM of MtNFP, where there is a single Gx_3_G motif, suggesting a possible role of the TM regions in initiating or determining the interactions of the full-length proteins.

The crystal structure of the ECD of AtSOBIR1 revealed the similarity between this region and other LRR-RLKs that are involved in binding peptides and other small ligands (Hohmann and Hothorn 2019). However, it is the interacting RLPs that bind ligands, and constitutively formed SOBIR1-RLP complexes seem to require a ligand-dependent interaction step with co-receptors like BAK1 or other SERKs (Gust and Felix 2014). The interaction observed between MtSOBIR1 and MtNFP in *N. benthamiana* leaves also suggests that a similar mechanism might exist, involving a constitutive, ligand-independent MtSOBIR1-MtNFP complex.

### Do MtNFP and MtSOBIR1 intervene together in the control of pathogen immunity?

Many studies in different species have clearly demonstrated the positive roles that SOBIR1 proteins play in plant immunity and plant development (Leslie *et al*., 2010; van der Burgh *et al*., 2019). For example, AtSOBIR1 acts as a counter player of BIR1 that is an immune suppressor (Gao et al., 2009). The predicted high structural similarity between MtSOBIR1 and orthologs in other plants and its ability to substitute for AtSOBIR1 in a defence response, is therefore suggestive of a conserved role of MtSOBIR1 in plant immunity and/or developmental processes. Furthermore, our data suggest that MtSOBIR1 can contribute to an immune-associated response to plant pathogens in roots. The question then is whether MtSOBIR1 and MtNFP intervene together for their roles in immunity against root pathogens like *A. euteiches*, especially given that SOBIR1 has only ever been reported to interact with RLKs/RLPs with LRR domains (Snoeck et al 2023).

Firstly, we confirmed that *MtSOBIR1* is well expressed in roots and root hairs, like *MtNFP*, which is compatible with a role of *MtSOBIR1* together with MtNFP in early root-pathogen interactions. Secondly, in terms of disease susceptibility in the presence of *A. euteiches*, we observed a phenotype for two *Mtsobir1* mutant alleles (*Mtsobir1-1 and 1-3*), but no change in the infection level of *A. euteiches* as measured by RT-qPCR. In contrast, the phenotypes of allelic *Mtnfp* mutants are clear for both symptom development and infection levels (Rey et al 2013), suggesting a more important role for MtNFP in interaction with *A. euteiches*. In other plants, loss of function *sobir1* mutants can result in phenotypes, especially with fungal pathogens (Zhang et al. 2013) (Zhou et al. 2019); (Seifbarghi et al. 2020), while over-expression of *SOBIR1* can activate cell death and defence responses (Gao et al 2009). We were unsuccessful in over-expressing *MtSOBIR1* to test for increased tolerance to *A. euteiches*. Lastly, we showed that three immunity-associated genes are not induced in *Mtsobir1* mutant roots exposed to *A. euteiches* for 24h. These genes are considered to be part of a general defence-type response because of their activation by several pathogens, including for example *Phymaotrichopsis omnivore* (Rey et al 2013), suggesting that MtSOBIR1 intervenes in a non-specific immunity response. MtNFP is also implicated in a non-specific immunity response, because *Mtnfp* mutants are more susceptible to several plant pathogens (*A. euteiches, Verticillium albo-atrum, Colletotrichum trifolii*a and *Phythophora palmivora*) (Gough and Jacquet 2013). However, the genes that are no longer up-regulated in *Mtsobir1* mutant roots, as well as numerous other genes with similar profiles of activation by plant pathogens, are still induced in *Mtnfp* mutant roots exposed to *A. euteiches* (Rey et al 2013). This suggests that MtSOBIR1 and MtNFP intervene in different mechanisms of pathogen-induced immunity, involving different receptor complexes.

#### Does MtSOBIR1 control the symbiotic suppression of immunity?

In symbiosis, plant immunity has to be tightly controlled, and the up regulation of defence gene expression in *Mtsobir1* mutant nodules was evidence of defects in the control of plant immunity. Nodule defence responses are suppressed by different pathways including those controlled by *nodule with activated defense 1 (Nad1), defective in nitrogen-fixation 2 (Dnf2)* and *cysteine-rich receptor-like kinase* (*SymCRK)* (Berrabah *et al*., 2014, 2018; Wang *et al*., 2016), leading to healthy infected and fixing nodules. At least 2 of the genes misregulated in *Mtsobir1-1* nodules formed by *S. medicae* WSM419 (PR10 and NDR1) are also misregulated in nodules of nodule immunity mutants (Wang et al 2016); (Gourion et al. 2015; Berrabah, Ratet, and Gourion 2015). By analysing a double *Mtsobir1/Mtnad1* mutant, our results indicate that *SOBIR1* is neither involved in a receptor complex activating defence responses in *nad1* nodules nor can substitute for *NAD1* for the suppression of these responses. However, *SOBIR1* may be required for other pathways involving other defence suppressors such as *SymCRK* or *DNF2* (Cao et al. 2017) or its role in regulating defence gene expression may define an additional defence-suppression pathway. Since environmental conditions can affect the immunity/senescence balance in nodules (Berrabah, Bourcy, Cayrel, et al. 2014) (Berrabah et al. 2023), it would be interesting to test R108 *Mtsobir1* mutants with *S. medicae* WSM419 in a range of growth conditions to compare nitrogenase activities and defence/senescence gene expression levels.

Another indication of a role of MtSOBIR1 in controlling symbiotic plant immunity, was the appearance of brown and necrotic epidermal cells in R108 *Mtsobir1* mutant roots in response to rhizobia. These cells resemble the necrotic cells associated with aborted rhizobial infection events in alfalfa (Vasse, de Billy, and Truchet 1993), and could be seen in some cases adjacent to aborted infection events in R108 *Mtsobir1* mutants. Vasse et al. considered these cells as a Hypersensitive Reaction (HR), more commonly observed in incompatible plant-pathogen interactions, and that they are part of a mechanism by which plants regulate nodulation. To our knowledge, such cells are not commonly seen in other *Medicago* spp., but it could be that the defence responses in epidermal cells associated with aborted infection events are better suppressed and/or do not produce visible symptoms in other contexts. The presence of necrotic epidermal cells in R108 *Mtsobir1* mutant roots, together with the good expression level of *MtSOBIR1* in *M. truncatula* epidermal roots cells, suggests a certain level of incompatibility due to *MtSOBIR1* loss of function. Moreover, this likely contributed to the strong defect in infection thread formation in R108 *Mtsobir1* mutants with *S. medicae* WSM419.

### Do MtNFP and MtSOBIR1 intervene together in symbiosis?

For the comparison between the symbiotic roles of MtNFP and MtSOBIR1, an argument in favour of MtSOBIR1 intervening with MtNFP is the similar gene expression patterns, with both *MtNFP* and *MtSOBIR1* showing expression in root hairs and similar zones of nodules. However, while *Mtnfp* loss of function mutants are completely deficient in nodulation with both *S. meliloti* 2011 and *S. medicae* WSM419, we only found a role of MtSOBIR1 in interaction with *S. medicae* WSM419. Nevertheless, *MtNFP* is involved in symbiont specificity (Bensmihen, de Billy, and Gough 2011) and both *MtNFP* and *MtSOBIR1* are involved in infection thread formation (Arrighi et al 2016 and this work). Indeed, a striking phenotype of R108 *Mtsobir1* mutants with *S. medicae* WSM419 was the reduced number of nodules, showing an effect on the early stages of the interaction in roots and root hairs. Except for the rare infection events that led to nodules, we observed that all rhizobial infection events were blocked at an early stage, before the formation of an infection thread. A complex mechanism controls the formation of infection threads, involving multiple plant and rhizobial components (Tsyganova, Brewin, and Tsyganov 2021). Among them are plant LysM-RLKs, and a notable difference recently reported between A17 and R108 is the presence of a LysM-RLK, MtLYK2bis, specifically in R108. This receptor confers on R108 the ability to nodulate with certain rhizobial strains, including a *S. meliloti* 2011 mutant producing modified Nod factors, and *S. medicae* WSM419 (Luu et al 2022). In addition, MtLYK2bis is an interactor of MtNFP. The necessity or not for MtLYK2bis is proposed to be linked to the production of differing proportions of non-*O*-acetylated Nod factors among rhizobial strains. There is no evidence that this is the case for *S. medicae* WSM419, but it is interesting that efficient nodulation in R108 by *S. medicae* WSM419 requires both MtLYK2bis and MtSOBIR1, both interactors of MtNFP. A *M. truncatula* LysM-RLK closely related to MtLYK2bis is MtLYK3, another MtNFP interactor that controls rhizobial infection in A17, but is dispensable in R108 (Catoira et al. 2001); ‘Moling et al 2014; Luu et al 2023). When MtLYK2bis or MtLYK3 is co-expressed with MtNFP in *N. benthiama* leaves, both induce cell death (Luu et al 2022); (Pietraszewska-Bogiel et al. 2013). MtSOBIR1 is likely to be able to trans-phosphorylate MtLYK2-bis, given that MtSOBIR1-KD could trans-phosphorylate other LysM-RLKs in addition to MtNFP, including the KD of MtLYK3. Given this and the observed defence gene induction in R108 *Mtsobir1* nodules, we could hypothesise that the control of defence reactions underlies, or at least partly explains, why the combined presence of MtLYK2bis and MtSOBIR1, together with MtNFP, allows efficient nodulation by *S. medicae* WSM419.

#### Why does MtSOBIR1 control specificity of nodulation?

The strong *Mtsobir1* phenotype depended on both the rhizobial strain *S. medicae* WSM419, and the R108 background of *M. truncatula*. *S. medicae* WSM419 is an acid tolerant bacterium and an efficient symbiont of a broad range of annual medics of Mediterranean origin (Garau et al. 2005). *S. medicae* WSM419 is notably a more efficient symbiont of A17 than *S. meliloti* 1021 (closely related to *S. meliloti* 2011) (Kazmierczak et al 2017). Between *S. medicae* WSM419 and *S. meliloti* 1021, there are no significant differences in the Nod factor-encoding *nod* genes (Baxter et al. 2021). Also in the *S. medicae* WSM419 – *S. meliloti* 1021 comparison, orthologous protein predictions show an enrichment in many GO terms for *S. medicae* WSM419-specific proteins, for example cell wall macromolecule catabolic process, peptidoglycan catabolic process and lysozyme activity, suggesting possible *S. medicae* WSM419 specificities for plant cell wall degradation (Baxter et al 2021). Proteomic studies have confirmed that *S. medicae* WSM419 produces many proteins that do not have a high level of similarity to any *S. meliloti* 1021 protein, and the transfer of three of these genes into *S. meliloti* 1021 improved plant growth on A17 (Yurgel et al. 2021) (Ghosh et al. 2021). Among these three genes, was an ACC-deaminase and a novel glucosidase, suggesting differences related to ethylene and exopolysaccharides (EPS), respectively, two components with known roles in nodulation. Moreover, we showed that the nodulation ability of *S. medicae* WSM419 on R108 *Mtsobir1-1* mutant plants was improved with the ethylene inhibitor AVG, while rhizobial surface EPS is involved in suppressing host defence (Jones et al. 2008). The partial improvement of *S. medicae* WSM419 nodulation of R108 *Mtsobir1* mutant plants conferred by the presence of AVG could be because of a high ethylene induction and/or by a high sensitivity to ethylene in the *S. medicae* WSM419 - R108 *Mtsobir1* combination.

Although generally known as *M. truncatula* subsp. *tricycla,* a recent plastid phylogenomics study has reported that the R108 accession may have originated from the *M. truncatula* sister subclade *M. littoralis* (Choi et al. 2022). Unsurprisingly, between the A17 and R108 genotypes, there are many differences, in addition to rhizobial strain preferences. For example, different sensitivities to salt stress and drought, different response and adaptation to iron deficiency, different responses to mineral toxicity of aluminium and sodium (Luo et al. 2016) (Li et al. 2014) (Wang et al. 2014) (de Lorenzo et al. 2007). These phenotypic variations could result from differences in the genomic structural variations between A17 and R108 (Li et al. 2022). These include variations within the nodule-specific cysteine-rich (NCR) gene family that encodes antimicrobial peptides essential for bacteroid differentiation and viability in nodules (Li et al 2022). Two such NCR genes, NFS1 and NFS2, control discrimination against strains of rhizobium in specific *M. truncatula* genotypes (Yang et al. 2017). Furthermore, big variations are detected in gene expression (for example, of NCR genes, defence genes and oxidation-reduction–related genes), due to the combined effects of both host plant and symbiont strain, with A17 showing less variation in response to symbiont identity than R108 (Burghardt et al. 2017). For the strain-dependent nodulation phenotype of the *M. truncatula efd* mutant, both the level and the timing of *NCR* gene expression are proposed to be involved (Jardinaud et al 2022). Other mechanisms of nodulation specificity also exist, for example, the *Rj4* gene in soybean that encodes a thaumatin-like protein (Tang et al. 2016). There are therefore many possibilities to explain the genotype- and strain-specific symbiotic phenotypes we found for *Mtsobir1* mutants.

Together, these data suggest that MtSOBIR1, like MtNFP, plays a dual role in symbiosis and plant immunity, and that MtSOBIR1 could control immunity in both pathogenic and beneficial situations, with positive or negative roles, respectively.

## Supporting information

Sarrette et al Supplementary data

## Acknowledgments

We are very grateful to Peter Kalo for providing *Mtnad1* mutant seeds. This work was funded by the Agence National de la Recherche (ANR-DUALITY project, ANR-20-CE20-0017-01) and by the Fédération de la Recherche (FR-AIB project CHAIN). This work was also supported in part by the Région Occitanie, European funds (Fonds Européens de DEveloppement Régional, FEDER), Toulouse Métropole, and by the French Ministry of Research with the Investissement d’Avenir Infrastructures Nationales en Biologie et Santé program (ProFI, Proteomics French Infrastructure project, ANR-10-INBS-08).This study is set in the framework of the “Laboratoire d’Excellences (LABEX) TULIP (ANR 10 LABX 41) and of the Ecole Universitaire de Recherche (EUR) TULIP GS (ANR 18 EURE 0019). The *Medicago truncatula Tnt1* mutant plants used in this research project, which are jointly owned by the Centre National de la Recherche Scientifique, were obtained from the Noble Research Institute, LLC and were created through research funded, in part, by a grant from National Science Foundation, NSF-0703285. TBL, BS and AJ gratefully acknowledge receipt of grants to fund their PhDs; from a Bourse d’Excellence from the Ambassade de France au Vietnam, from the Ministère de l’Enseignement Supérieur et de la Recherche, France, and from both the SPE department of INRAE and the Occitanie Région in France, respectively.

## Supplementary data

**Table S1. List of primers used in this study**

**Figure S1. Verification of the interaction between the MtNFP kinase domain and an MtSOBIR1 clone on yeast selective medium.**

**Figure S2. MtSOBIR1 shows high homology with AtSOBIR1. An Amino acid alignment of SOBIR1 in *Arabidopsis thaliana* and *Medicago truncatula*, with conserved features highlighted.**

**Figure S3. Phylogenetic tree of MtSOBIR1 with AtSOBIR1 and homologs in other leguminous species.**

**Figure S4: MtSOBIR1 complements *N. benthamiana sobir1*.**

**Figure S5. Transphosphorylation by MtSOBIR1 of NFP-KD and KDs of related proteins.**

**Figure S6. Label-free visualization of autophosphorylated MtSOBIR1-KD and trans-phosphorylated NFP for mass spectrometry analysis.**

**Table S2. Autophosphorylation sites of MtSOBIR1-KD**

**Figure S7 Autophosphosites of MtSOBIR1-KD identified in this paper in comparison with AtSOBIR1-KD sites.**

**Table S3. NFP-KD is transphosphorylated by MtSOBIR1-KD on 9 Serine/Threonine residues.**

**Figure S8. Alignment of the kinase domains of MtNFP and LjNFR5 showing transphosphorylated sites.**

**Figure S9. Analyses of NFP-KD phosphorylation by MtSOBIR1-KD**

**Figure S10. Expression pattern of *MtSOBIR1* in *Medicago truncatula* roots, root hairs and nodules in comparison with *MtNFP*.**

**Figure S11. Expression pattern of *MtSOBIR1* in *Medicago truncatula* roots in response to *Aphanomyces euteiches* in comparison with *MtNFP***

**Figure S12. *Mtsobir1* mutants and *Mtsobir1-1* expression analysis.**

**Figure S13. *Mtsobir1-3* and *Mtsobir1-CRP* mutant phenotypes in response to Aphanomyces euteiches.**

**Figure S14. Nodulation phenotypes of *Mtsobir1* mutants with *Sinorhizobium meliloti* 2011.**

**Figure S15. Infection and kinetics of nodulation on *Mtsobir1-1* mutant and WT plants with *S. meliloti 2011* in *in vitro* conditions.**

**Figure S16. *MtSOBIR1* does not intervene in the activation of defence responses that are suppressed by NAD1**

**Figure S17. Kinetics of nodulation tests on *Mtsobir1-1* mutant and WT plants with *S. medicae* WSM419 in *in vitro* conditions**

**Figure S18. Nodulation tests on *Mtsobir1-1* mutant and WT plants with *S. medicae* WSM419 in *in vitro* conditions with/without 100 mM AVG**

**Figure S19. Nodulation and nitrogenase activity of *Mtsobir1-1* plants inoculated by *S. meliloti 2011* or *S. medicae* WSM419.**

**Figure S20. Brown epidermal cells observed for *Mtsobir1-1* and *Mtsobir1-3* mutants when inoculated in *in vitro* conditions.**

