## Supplementary material for "*Medicago truncatula* SOBIR1 controls specificity in the Rhizobium-legume symbiosis": Sarrette et al Supplementary data

| Primer_ID | Sequence | Gene (Mt V5) | Purposes |
|---|---|---|---|
| NEP013_F | ATGGCTAAGAACAACAACATC | MtrunA17_Chr3g0116591 | full *SOBIR1* coding sequence |
| NEP013_R | GTGCTTGATTTGGTACAACAT | MtrunA17_Chr3g0116591 | |
| SOBcrpF1 | ACTACAAACAACCTATGTAACA | MtrunA17_Chr3g0116591 | Sequencing CRISP/Cas9 plants |
| SOBcrpR1 | TAAATCCTCAGCTTTCTTAATCA | MtrunA17_Chr3g0116591 | |
| proSOBIR1_GG_F1 | GGGTCTCCAAATCCAAGGATATTGGGCAAAGA | MtrunA17_Chr3g0116591 | Cloning pSobir1 promoter |
| proSOBIR1_GG_R1 | AAACAGAACACAAACTTGGCCTCACAAATGAGACCT | MtrunA17_Chr3g0116591 | |
| SymCRK_F | GATTTCTGTGTTGAAGCTTGGCT | MtrunA17_Chr3g0119041 | qRT-PCR |
| SymCRK_R | ACATCAGAAGTGAACTCTCTGCAA | | |
| CHS_F | TTGCGCAGTGTCGCTATATG | MtrunA17_Chr3g0119041 | qRT-PCR |
| CHS_F | TGAGGGTACATGTTGAACACTAGA | | |
| PR 10.2_F1 | AGCGAAATTGGTTGAAGGC | MtrunA17_Chr4g0067951 | qRT-PCR |
| PR 10.2_R1 | TCTCCTTTGGTTTGGTATTTAACT | | |
| Thaumatin_F | GGCGCAATCCCACCAGCAAC | MtrunA17_Chr1g0180221 | qRT-PCR |
| Thaumatin_R | ACCACTCCCACCTTGTGGCG | | |
| Prx67_F | TTGCTAGACGACACCTCCAC | MtrunA17_Chr4g0043391 | qRT-PCR |
| Prx67_R | CGAGCTGCTACTGCTACGAT | | |
| MtRbohD_F | GAGCACCGGCACCTATTAAA | MtrunA17_Chr3g0131361 | qRT-PCR |
| MtRbohD_R | CGTCTTTGTTCCCGTCCTT | | |
| MtBGL_F | CAAATTGGGTCCAAAAATATGTGAC | MtrunA17_Chr4g0038981 | qRT-PCR |
| MtBGL_R | GCACCATCATTGGGTGGATATGAAG | | |
| NDR1_F | AACAACAACACCCTCCTCCA | MtrunA17_Chr5g0433201 | qRT-PCR |
| NDR1_R | TTGAATGTGACTGCCAAACTG | | |
| VSP_F | GACCTTTGGGTGTTTGACATTGA | MtrunA17_Chr8g0340821 | qRT-PCR |
| VSP_R | TCCTTCTGTTTGAGTGGTCTTCCT | | |
| EF1A_F | TAACAAGATGGATGCTACCACCC | MtrunA17_Chr6g0458091 | qRT-PCR |
| EF1A_R | GATTTCATCGTACCTAGCCTTTGA | | |
| A38_F | TCGTGGTGGTGGTTATCAAA | MtrunA17_Chr4g0061551 | qRT-PCR |
| A38_R | TTCAGACCTTCCCATTGACA | | |
| Ae_TUB_F | CGGCTCTGGTTTGGGTAGTTT | *Aphanomyces euteiches* | qRT-PCR |
| Ae_TUB_R | AACCGAGCTTGCTCTTGCG | *Aphanomyces euteiches* | qRT-PCR |
| NFPbs_F | ACTTCATCGTCCGAGACTGC | MtrunA17_Chr5g0403371 | qRT-PCR |
| NFPbs_R | GCAGACCGATTGCAACATCC | MtrunA17_Chr5g0403371 | qRT-PCR |
| NEP018F7 | CGGAGATTGACACAGTCGGT | MtrunA17_Chr3g0116591 | qRT-PCR *sobir1_3* |
| NEP018R7 | GCCAACCAATCCAACTCTGC | MtrunA17_Chr3g0116591 | qRT-PCR *sobir1_3* |
| NEP018F6 | ACACAAACTTGGCCTCAATGG | MtrunA17_Chr3g0116591 | qRT-PCR *sobir1_1* |
| NEP018R6 | TGTCTGAAGGGTGAAGGGTG | MtrunA17_Chr3g0116591 | qRT-PCR *sobir1_1* |

**Table S1** List of primers used in this study

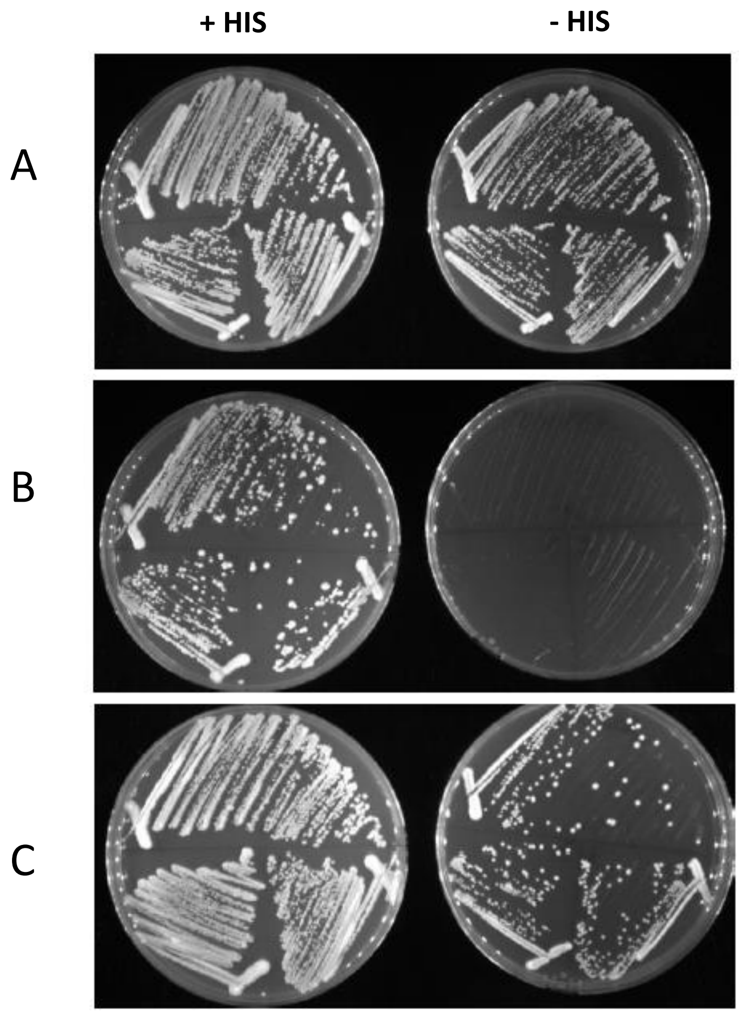


**Figure S1. Verification of the interaction between the MtNFP kinase domain and an MtSOBIR1 clone on yeast selective medium.** A, Hybrigenics positive controls (Bait : SMAD; Prey SMURF); B, Bait MtNFP; Prey : empty vector pB27; C, Bait MtNFP; prey : MtSOBIR1

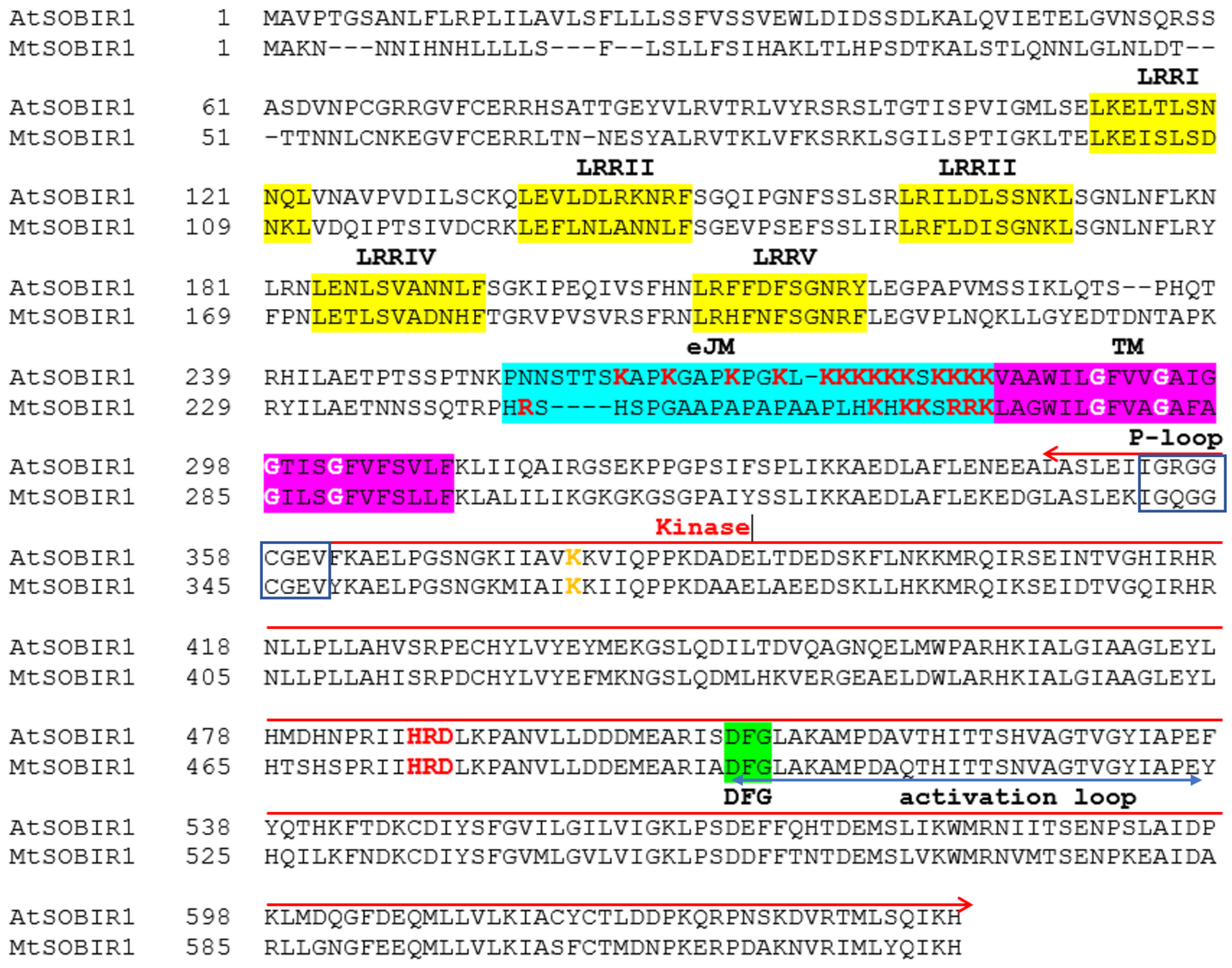


**Figure S2. MtSOBIR1 shows high homology with AtSOBIR1**. An Amino acid alignment of SOBIR1 in *Arabidopsis thaliana* and *Medicago truncatula*, with conserved features highlighted. Yellow: the LRR motifs; blue: the extracellular JuxtaMembrane (eJM) region; pink: the transmembrane (TM) region. Typical features of active kinase domains (P-loop, HRD and DRF motifs, the activation loop) are indicated.

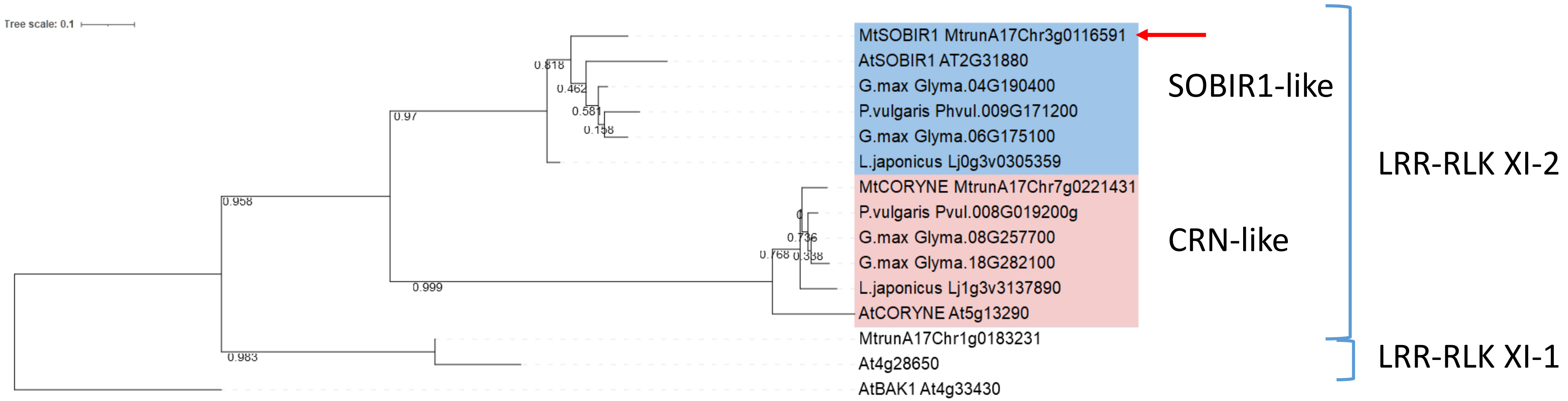


**Figure S3. Phylogenetic tree of MtSOBIR1 with homologs in other leguminous species and *Arabidopsis thaliana*, as well as closely related LRR-RLK proteins in the XI-1 and XI-2 subgroups, and with AtBAK1 as an outgroup.** This maximum likelihood tree, including bootstrap values, was made from a multiple sequence alignment of SOBIR1 homologues, using phylogeny.fr and default settings. MtSOBIR1 is indicated with a red arrow.

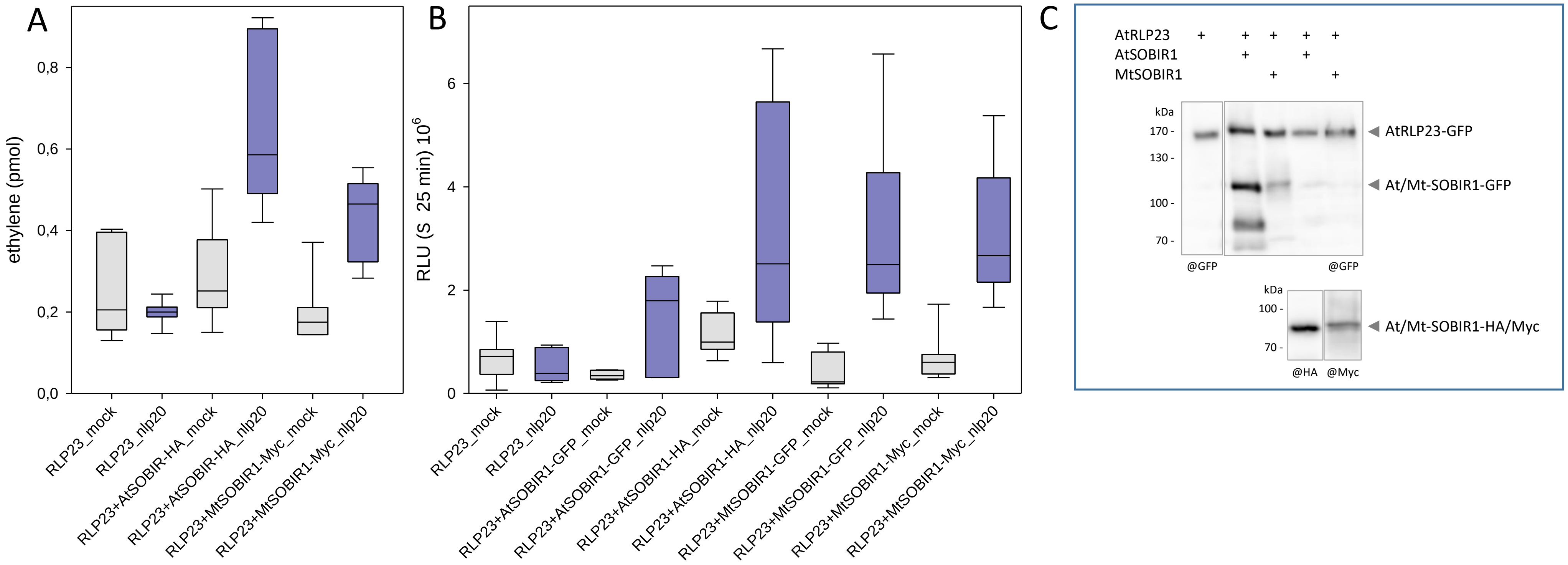


**Figure S4: MtSOBIR1 complements *Nicotiana benthamiana sobir1*.** SOBIR1 from *M. truncatula* or *A. thaliana* were transiently expressed in CRISPR/CAS mutants of *Nb*SOBIR1 (1) in combination with AtRLP23. A, Induction of ethylene synthesis in leaf pieces expressing AtRLP23 only, or AtRLP23 and Mt- or AtSOBIR1 after mock treatment or treatment with 100 nM nlp20 for 3,5 h. B, ROS production measured as relative light units (RLU, integral over 25 min) after mock treatment or treatment with 100 nM nlp20. Data are from 2 independent experiments with 3 or 4 technical replicates each. C, Western blot to verify the expression of the different receptors.

1. Huang et al. (2021) Knocking out *SOBIR1* in *Nicotiana benthamiana* abolishes functionality of transgenic receptor-like protein Cf-4. Plant Physiol. 185:290-294

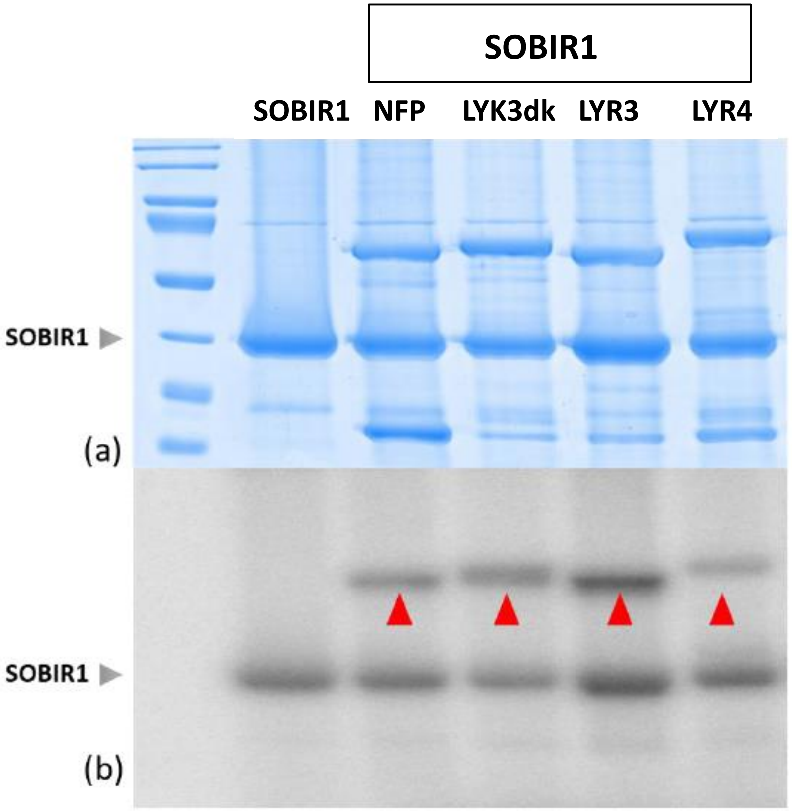


**Figure S5. Transphosphorylation by MtSOBIR1 of NFP-KD and KDs of related proteins**. Soluble 6His-GST-cleaved SOBIR1-KD was incubated alone (lane 1) or with purified GST-NFP-KD, GST-LYK3deadKD, GST-LYR3-KD and GST-LYR4-KD (lanes 2-5) in the presence of radioactive [ɣ-32P] ATP at 25ºC for 1h. Assays were analysed by SDS/PAGE, followed by a coomassie staining (a) and phosphor imaging (b). Transphosphorylated proteins are marked by red arrowheads on the phosphor-image.

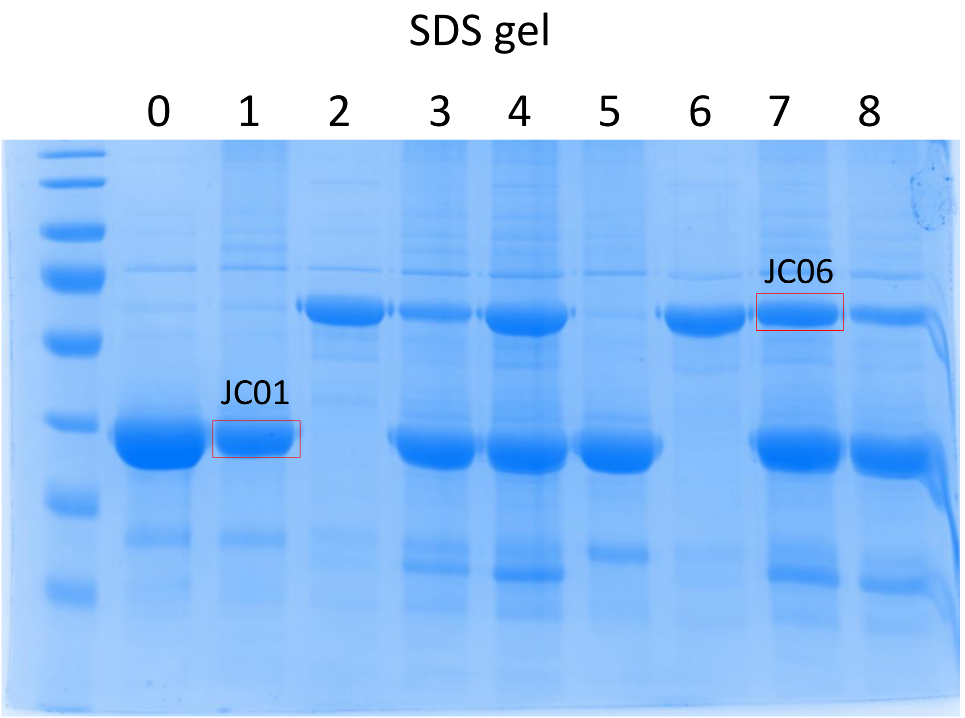


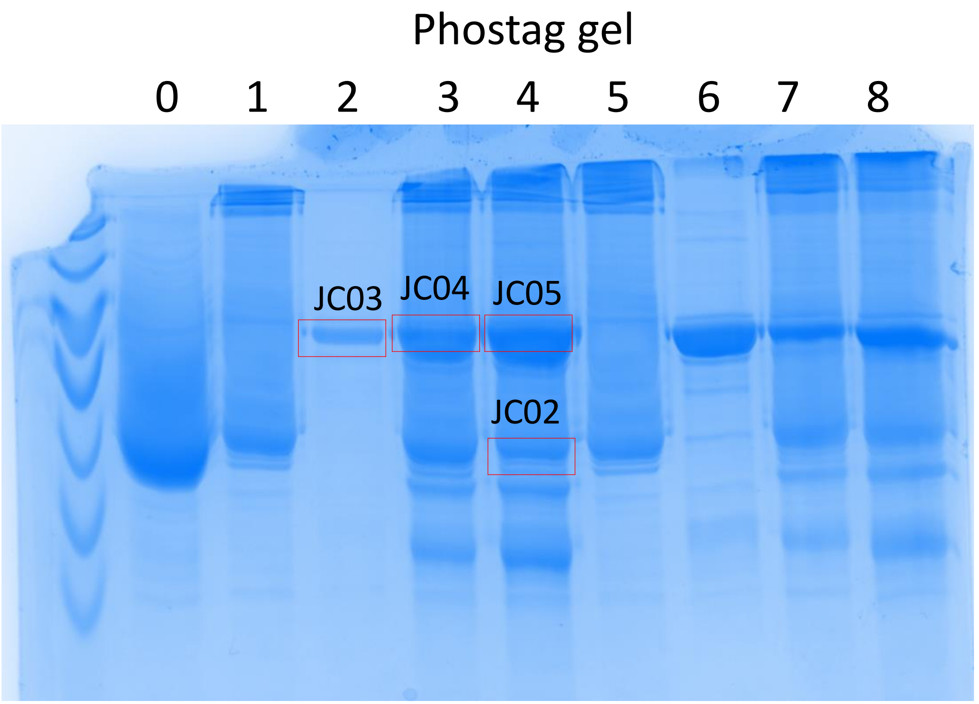


| No | P1 | P2 | |
|---|---|---|---|
| 0 | SOBIR1-KD | | no ATP |
| 1 | SOBIR1-KD | | |
| 2 | 6His-GST-NFP-KD | | |
| 3 | SOBIR1-KD | 6His-GST-NFP-KD | |
| 4 | SOBIR1-KD | 6His-GST-NFP-KD | |
| 5 | SOBIR1-KD | | |
| 6 | 6His-GST-NFP-KD | | |
| 7 | SOBIR1-KD | 6His-GST-NFP-KD | |
| 8 | SOBIR1-KD | 6His-GST-NFP-KD | |

Samples in red boxes were collected for proteomic study:
JC01: Phosphorylated MtSOBIR1-KD alone
JC02: Phosphorylated MtSOBIR1-KD with NFP-KD
JC03: NFP-KD alone
JC04, JC05, JC06: Phosphorylated NFP-KD by MtSOBIR1-KD

**Figure S6. Label-free visualization of autophosphorylated MtSOBIR1-KD and trans-phosphorylated MtNFP-KD for mass spectrometry analysis.** Proteins were analysed by SDS-PAGE, followed by a Coomassie staining.

| Peptide | Calculated_mass | Charge | Expt. Moz | Delta_moz | Site | Other modifications |
|---|---|---|---|---|---|---|
| 307-GKGSGPAIYSSLIK-320 | 1456.732666 | 2 | 729.3725 | -0.001109 | S310 | |
| 307-GKGSGPAIYSSLIK-320 | 1456.732666 | 2 | 729.3726 | -0.001009 | S317 | |
| 307-GKGSGPAIYSSLIK-320 | 1456.732666 | 2 | 729.3727 | -0.000909 | S316 | |
| 309-GSGPAIYSSLIKK-321 | 1479.677521 | 2 | 740.8446 | -0.001436 | Y315 and S316 or S317 | |
| 321-KAEDLAFLEKEDGLASLEK-339 | 2185.055481 | 2 | 1093.5363 | 0.001283 | S336 | |
| 340-IGQGGCGEVYKAELPGSNGK-359 | 2099.934708 | 3 | 700.9832 | -0.002312 | S356 | Carbamidomethyl (C345) |
| 366-IIQPPKDAAELAEEDSKLLHK-386 | 2424.230087 | 2 | 1213.1227 | 0.00038 | S381 | |
| 390-QIKSEIDTVGQIR-402 | 1565.781418 | 2 | 783.8965 | -0.001485 | S393 | |
| 390-QIKSEIDTVGQIR-402 | 1645.747742 | 2 | 823.8794 | -0.001747 | S393 and T397 | |
| 405-NLLPLLAHISRPDCHYLVYEFMK-427 | 2908.416885 | 4 | 728.1102 | -0.001297 | S414 | Carbamidomethyl (C419) |
| 428-NGSLQDMLHK-437 | 1221.521301 | 2 | 611.7653 | -0.002627 | S430 | |
| 453-IALGIAAGLEYLHTSHSPR-471 | 2165.007126 | 3 | 722.6774 | 0.001082 | Y463 and S469 | |
| 453-IALGIAAGLEYLHTSHSPR-471 | 2085.040802 | 2 | 1043.5298 | 0.002123 | T466 | |
| 500-AMPDAQTHITTSNVAGTVGYIAPEYHQILK-529 | 3305.579163 | 3 | 1102.8679 | 0.000903 | S511 | |
| 500-AMPDAQTHITTSNVAGTVGYIAPEYHQILK-529 | 3305.579163 | 3 | 1102.8679 | 0.000903 | T509 | |
| 500-AMPDAQTHITTSNVAGTVGYIAPEYHQILK-529 | 3305.579163 | 3 | 1102.8679 | 0.000903 | T510 | |
| 500-AMPDAQTHITTSNVAGTVGYIAPEYHQILK-529 | 3305.579163 | 3 | 1102.8682 | 0.001203 | T506 | |
| 500-AMPDAQTHITTSNVAGTVGYIAPEYHQILK-529 | 3305.579163 | 3 | 1102.8686 | 0.001603 | T516 | |
| 551-LPSDDFFTNTDEMSLVK-567 | 2037.864243 | 3 | 680.2951 | -0.000257 | S563 | |
| 551-LPSDDFFTNTDEMSLVK-567 | 2037.864243 | 2 | 1019.9414 | 0.002002 | T559 | |
| 551-LPSDDFFTNTDEMSLVK-567 | 2037.864243 | 2 | 1019.9408 | 0.001402 | T557 | Oxidation (M562) |
| 571-NVMTSENPK-579 | 1098.44165 | 2 | 550.2288 | 0.000699 | T574 | |
| 571-NVMTSENPK-579 | 1114.436569 | 2 | 558.2287 | 0.00314 | S575 | Oxidation (M573) |
| 601-IASFCTMDNPK-611 | 1362.534912 | 2 | 682.274 | -0.000732 | T606 | Carbamidomethyl (C605) |

**Table S2. Autophosphorylation sites of MtSOBIR1-KD**

```
AtSOBIR1kin SEKPPGPSIF SPLIKKAEDL AFLENEEALA SLEIIGRGGC GEVFKAELPG
MtSOBIR1kin KGKGSGPAIY SSLIKKAEDL AFLEKEDGLA SLEKIGQGGC GEVYKAELPG

AtSOBIR1kin SNGKIIAVKK VIQPPKDADE LTDEDSKFLN KKMRQIRSEI NTVGHIRHRN
MtSOBIR1kin SNGKMIAIKK IIQPPKDAAE LAEEDSKLLH KKMRQIKSEI DTVGQIRHRN

AtSOBIR1kin LLPLLAHVSR PECHYLVYEY MEKGSLQDIL TDVQAGNQEL MWPARHKIAL
MtSOBIR1kin LLPLLAHISR PDCHYLVYEF MKNGSLQDML HKVERGEAEL DWLARHKIAL

AtSOBIR1kin GIAAGLEYLH MDHNPRIIHR DLKPANVLLD DDMEARISDF GLAKAMPDAV
MtSOBIR1kin GIAAGLEYLH TSHSPRIIHR DLKPANVLLD DEMEARIADF GLAKAMPDAQ

AtSOBIR1kin THITTSHVAG TVGYIAPEFY QTHKFTDKCD IYSFGVILGI LVIGKLPSDE
MtSOBIR1kin THITTSNVAG TVGYIAPEYH QILKFNDKCD IYSFGVMLGV LVIGKLPSDD

AtSOBIR1kin FFQHTDEMSL IKWMRNIITS ENPSLAIDPK LMDQGFDEQM LLVLKIACYC
MtSOBIR1kin FFTNTDEMSL VKWMRNVMTS ENPKEAIDAR LLGNGFEEQM LLVLKIASFC

AtSOBIR1kin TLDDPKQRPN SKDVRTMLSQ IKH
MtSOBIR1kin TMDNPKERPD AKNVRIMLYQ IKH
```

**Figure S7. Autophosphosites of MtSOBIR1-KD identified in this paper in comparison with AtSOBIR1-KD sites** (extracted from 1).

1. Mitra *et al.* (2015). An autophosphorylation site database for leucine-rich repeat receptor-like kinases in Arabidopsis thaliana. Plant J. 82: 1042-1060.

| Peptide | Calculated mass | Charge | Expt. Moz | Delta_moz | Site | Frequency |
|---|---|---|---|---|---|---|
| 322-IGE**S**VYK-328 | 874.383728 | 2 | 438.1995 | 0.00036 | **S326** | **2** |
| 397-T**S**NSVVSLTWSQR-409 | 1543.703186 | 2 | 772.8558 | -0.003069 | **S398** | **3** |
| 397-TSN**S**VVSLTWSQR-409 | 1543.703186 | 2 | 772.8577 | -0.001169 | **S400** | **3** |
| 397-TSNSVV**S**LTWSQR-409 | 1543.703186 | 3 | 515.5757 | 0.000695 | **S403** | **1** |
| 397-**T**SNSVVSLTWSQR-409 | 1543.703186 | 2 | 772.8558 | -0.003069 | **T397** | **2** |
| 575-SLTSGLDAEA**T**HVVTSVIAR-595 | 2106.035812 | 2 | 1054.0265 | 0.001318 | **T585** | **3** |
| 575-SLTSGLDAEATHVV**T**SVIAR-595 | 2106.035812 | 3 | 703.0191 | -0.000113 | **T590** | **2** |
| 575-SL**T**SGLDAEATHVVTSVIAR-595 | 2106.035812 | 2 | 1054.026 | 0.000818 | **T577** | **1** |
| 575-**S**LTSGLDAEATHVVTSVIAR-595 | 2106.035812 | 2 | 1054.026 | 0.000818 | **S575** | **2** |

**Table S3. NFP-KD is transphosphorylated by MtSOBIR1-KD on 9 Serine/Threonine residues.**
Identification of phosphosites of NFP-KD by MtSOBIR1-KD. The transphosphorylated NFP-KD was purified from acrylamide gel and analyzed by LC-MS/MS. The sites are highlighted in bold red on petides of the NFP-KD. Frequency indicates the number of samples in which the peptides are found.

```
                          *
LjNFR5  .......RRK KALNRTASSA ETADKLLSGV SGYVSKPNVY EIDEIMEATK
MtNFP   GPLGSPEFKM KRLNRSTSSS ETADKLLSGV SGYVSKPTMY EIDAIMEGTT

LjNFR5  DFSDECKVGE SVYKANIEGR VVAVKKIKEG GANEELKILQ KVNHGNLVKL
MtNFP   NLSDNCKIGE SVYKANIDGR VLAVKKIKKD .ASEELKILQ KVNHGNLVKL

LjNFR5  MGVSSGYDGN CFLVYEYAEN GSLAEWLFSK SSGTPNS... LTWSQRISIA
MtNFP   MGVSSDNDGN CFLVYEYAEN GSLEEWLFSE SSKTSNSVVS LTWSQRITIA

LjNFR5  VDVAVGLQYM HEHTYPRIIH RDITTSNILL DSNFKAKIAN FAMARTSTNP
MtNFP   MDVAIGLQYM HEHTYPRIIH RDITTSNILL GSNFKAKIAN FGMARTSTNS

LjNFR5  MMPKIDVFAF GVLLIELLTG RKAMTTKENG EVVMLWKDMW EIFDIEENRE
MtNFP   MMPKIDVFAF GVVLIELLTG KKAMTTKENG EVVILWKDFW KIFDLEGNRE

LjNFR5  ERIRKWMDPN LESFYHIDNA LSLASLAVNC TADKSLSRPS MAEIVLSLSF
MtNFP   ERLRKWMDPK LESFYPIDNA LSLASLAVNC TADKSLSRPT IAEIVLCLSL

LjNFR5  LTQQSSNPTL ERSLTSSGLD VEDDAHIVTS ITAR
MtNFP   LNQPSSEPML ERSLTS.GLD AEA.THVVTS VIAR
```

**Figure S8. Alignment of the kinase domains of MtNFP and LjNFR5 showing sites transphosphorylated on MtNFP-KD by MtSOBIR1 (in red) and sites transphosphorylated on LjNFR5-KD by LjNFR1-KD and LjSYMRK-KD (in blue).** The asterisk corresponds to the only site in LjNFR5-KD phosphoryated by LjNFR1-KD. This site and the 7 other blue sites of LjNFR5-KD are phosphorylated by LjSYMRK-KD. The site highlighted in yellow corresponds to the only phosphorylated site that is conserved between MtNFP-KD transphosphorylated by MtSOBIR1-KD, and LjNFR5-KD transphosphorylated by LjSYMRK-KD (1).


1. Madsen *et al*. (2011). Autophosphorylation is essential for the in vivo function of the Lotus japonicus Nod factor receptor 1 and receptor-mediated signaling in cooperation with Nod factor receptor 5. Plant J. 65:404-417.

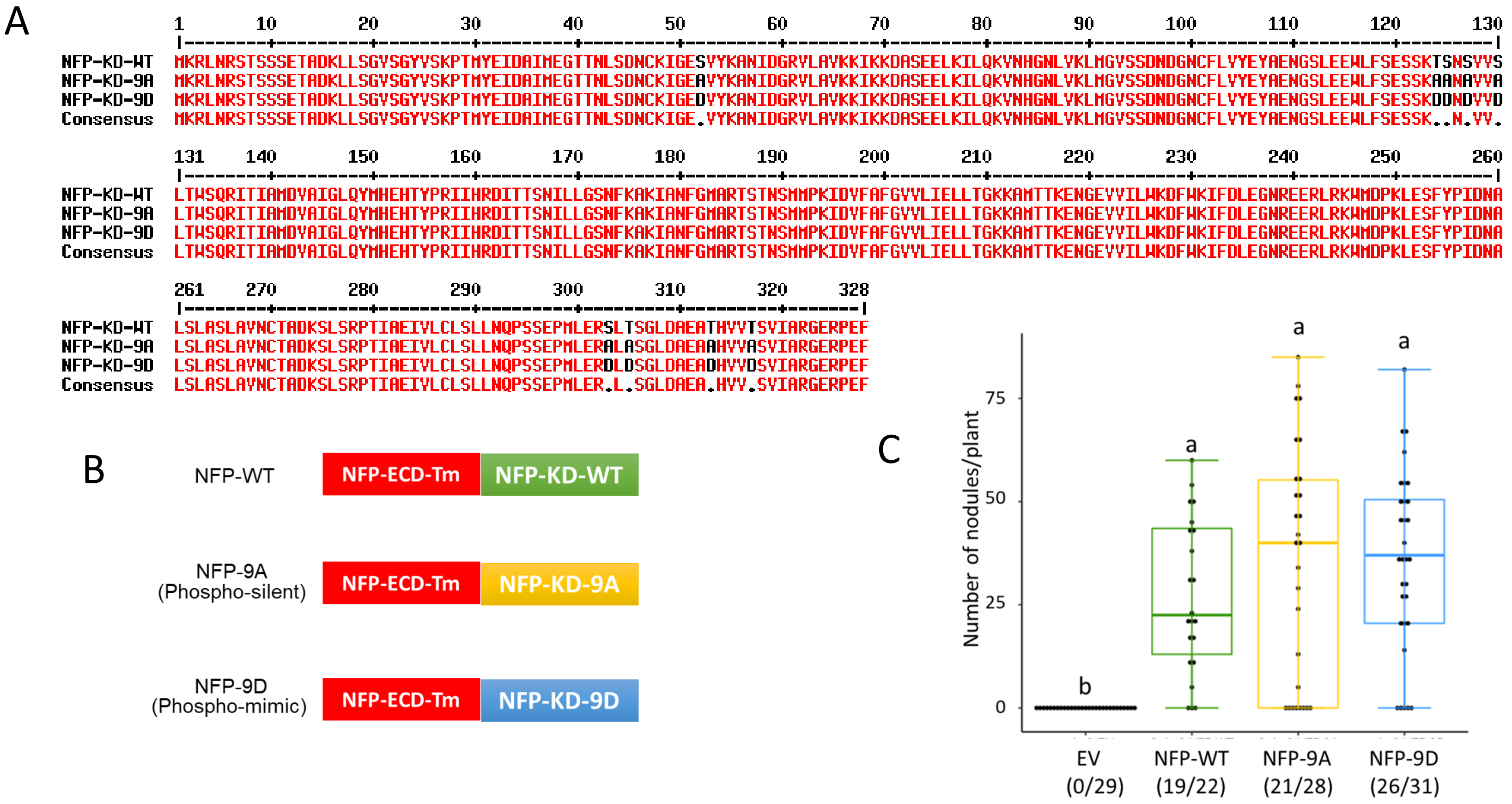


**Figure S9. Analyses of NFP-KD phosphorylation by MtSOBIR1-KD.** (a) Amino acid alignment of WT NFP-KD and the two phospho-versions of NFP-KD generated for the complementation study. In the phospho-silent (NFP-9A), all 9 detected sites were mutated to Alanine while in phosphor-mimic (NFP-9D), those sites were mutated to Aspatate (b) Schematic representation of the constructs used for complementation assays. (c) Complementation of the Mt*nfp-2* mutant with different phospho-versions of NFP-KD. *nfp-2* mutant roots were transformed with constructs of EV, NFP-WT, NFP-9A and NFP-9D using *A. rhizogenes*. Transformed plants were analysed at 3wpi. Statistical analyses were performed on the number of nodules using ANOVA test ($P < 0.05$). Lowercase letters indicate statistical significance. Numbers below indicate number of nodulated plants/total transformed plants.

MtSOBIR1

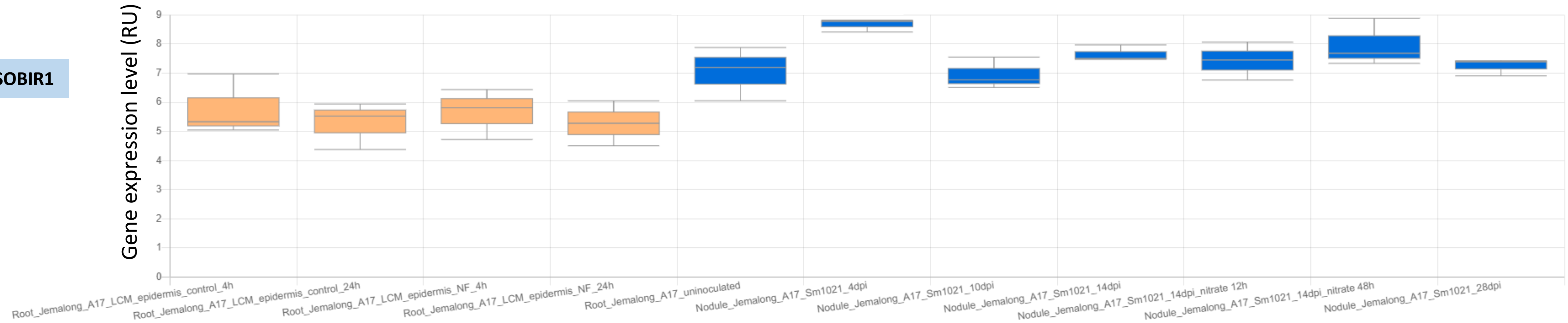


MtNFP

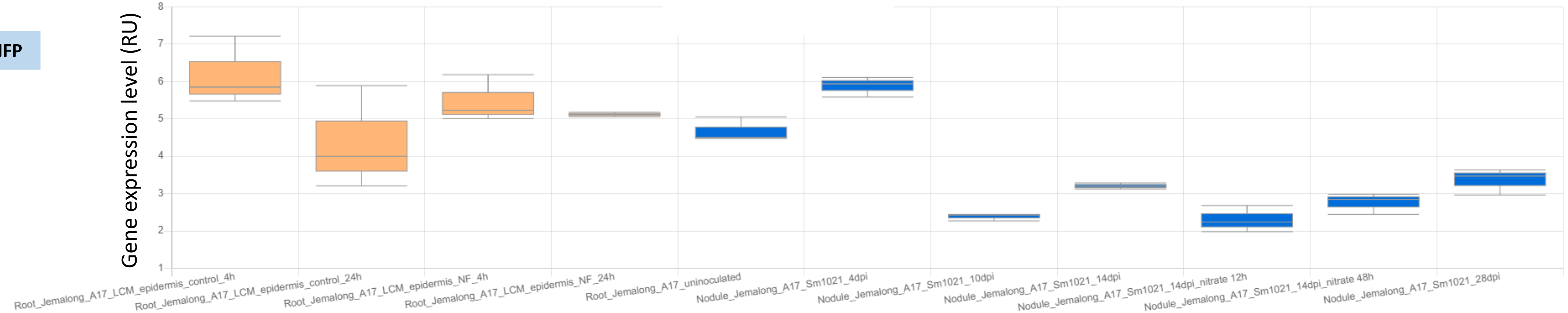


**Figure S10. Published expression data from MtExpressV3 (https://lipm-browsers.toulouse.inra.fr/pub/expressionAtlas/app/v3/) (1) for MtSOBIR1 (MtrunA17_Chr3g0116591) and MtNFP (MtrunA17_Chr5g0403371), and corresponding to symbiotic conditions from the indicated references.**

Jardinaud et al (2016) A Laser Dissection-RNAseq Analysis Highlights the Activation of Cytokinin Pathways by Nod Factors in the Medicago truncatula Root Epidermis. Plant Phys. 171:2256-76

De Bang et al (2017) Genome-wide Analysis of Small Signaling Peptides in Medicago truncatula with an Emphasis on Macro-nutrient Regulation of Root and Nodule Development. Plant Phys. 175:1669-89

1. Carrere et al. (2021). MtExpress, a comprehensive and curated RNAseq-based gene expression atlas for the model legume *Medicago truncatula*. *Plant Cell Physiol* 62. 1494-1500.

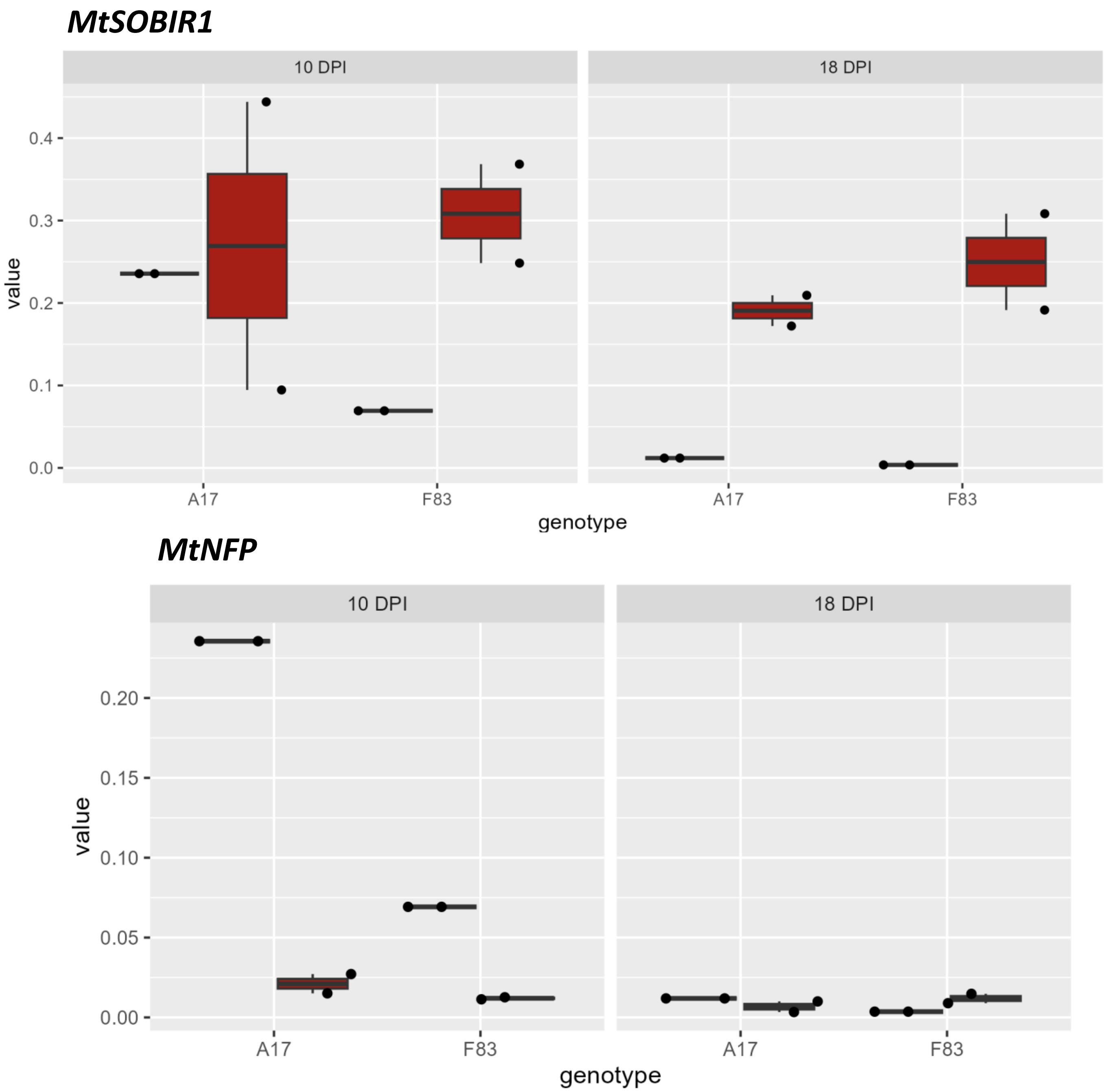


**Figure S11. Expression pattern of *MtSOBIR1* and *MtNFP* in *Medicago truncatula* roots in response to *Aphanomyces euteiches.*** RT-q-PCR analysis of *MtSOBIR1* and *MtNFP* expression in roots of *M. truncatula* A17 (tolerant) and F83 (susceptible) at 10 and 18 dpi with *A. euteiches*. Blue = non-inoculated; red = inoculated.

A

B

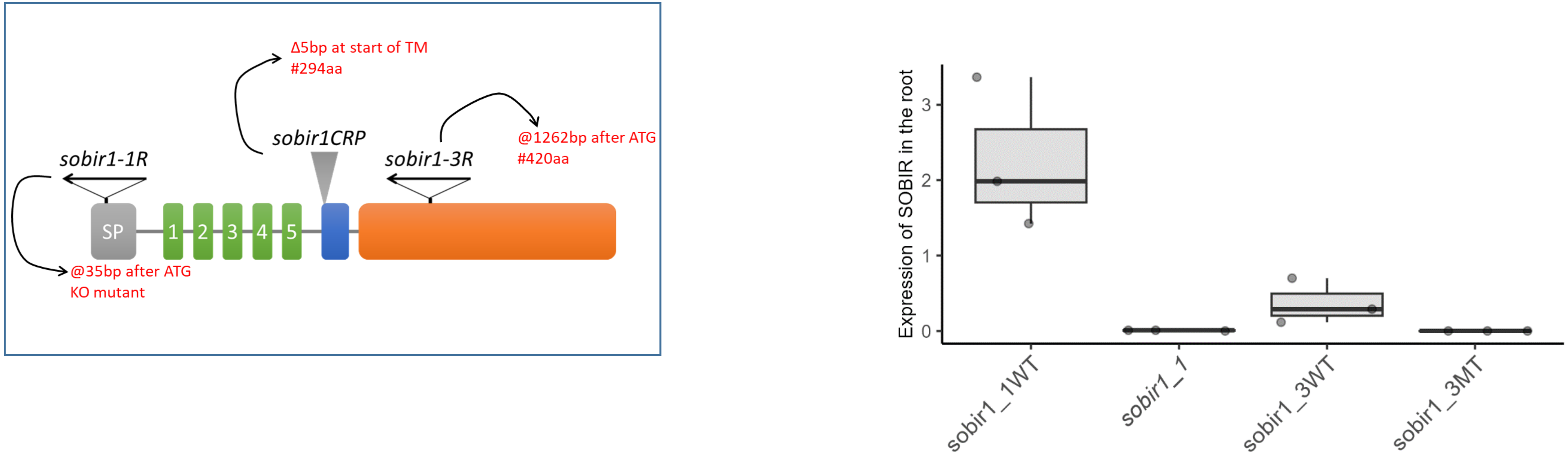


**Figure S12. *Mtsobir1* mutants and *Mtsobir1-1* expression analysis.** A, Schematic representation of the MtSOBIR1 protein and positions of the mutations used in this study. *MtSobir1-1* and *Mtsobir1-3* are Tnt1 insertion mutants, while *Mtsobir1CRP* is a CrispRCas9 mutant. B, Expression analysis by RT-qPCR of *MtSOBIR1* in the *Mtsobir1-1* and *Mtsobir1-3* mutants and their WT control lines.

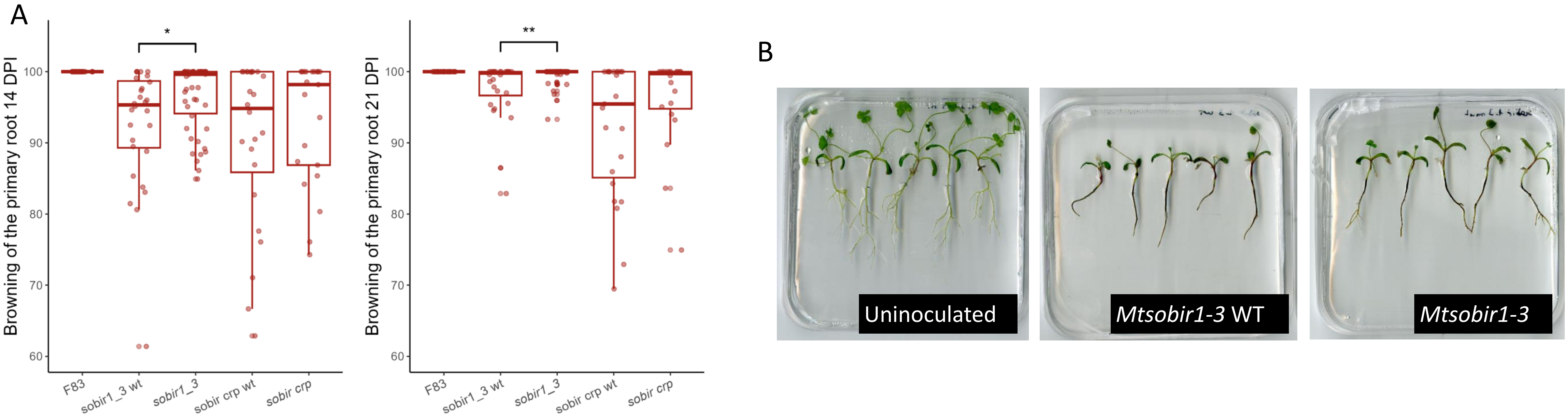


**Figure S13. *Mtsobir1-3* and *Mtsobir1-CRP* mutant phenotypes in response to *Aphanomyces euteiches.*** A, Symptom development at 14 and 21 dpi in *M. truncatula* F83 (susceptible), *Mtsobir1-3* WT*, Mtsobir1-3* mutant*, Mtsobir1-CRP WT* and *Mtsobir1-CRP* mutant following *A. euteiches* inoculation. B, Photos of seedlings of *Mtsobir1-3,* and control plants 21 dpi following *A. euteiches* inoculation.

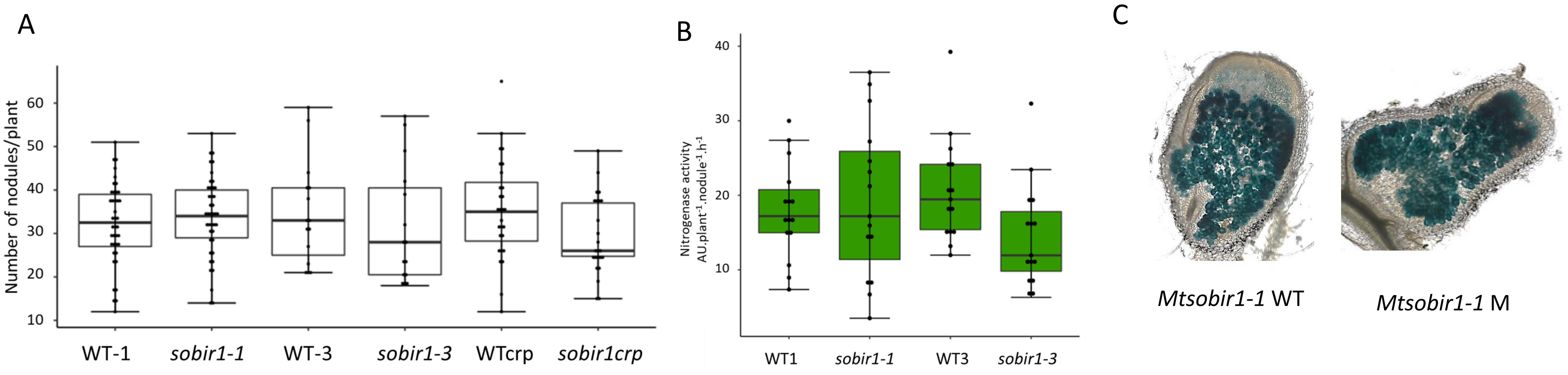


**Figure S14. Nodulation phenotypes of *Mtsobir1* mutants with *Sinorhizobium meliloti* 2011**. A, Nodulation tests in pot-grown plants with the three mutant alleles of *MtSOBIR1* and *S. meliloti* 2011. R108, *sobir1-1* WTL, *sobir1-3* WTL, and WTcrp were used as controls. The total numbers of nodules were scored at 21dpi. B, Nitrogenase activity, analysed by the acetylene reduction assay, at 21 dpi for *MtSobir1-1* and *Mtsobir1-3* plants and their WT controls, all nodulated by *S. meliloti* 2011. C, Nodule sections of *MtSobir1-1* mutant and its WT control, showing *S. meliloti* 2011 rhizobial colonization in blue (rhizobia constitutively expressing a LacZ fusion). Statistical analyses were performed using one-way ANOVA, Turkey ($P < 0.05$), and no statistically significant differences were observed between samples.

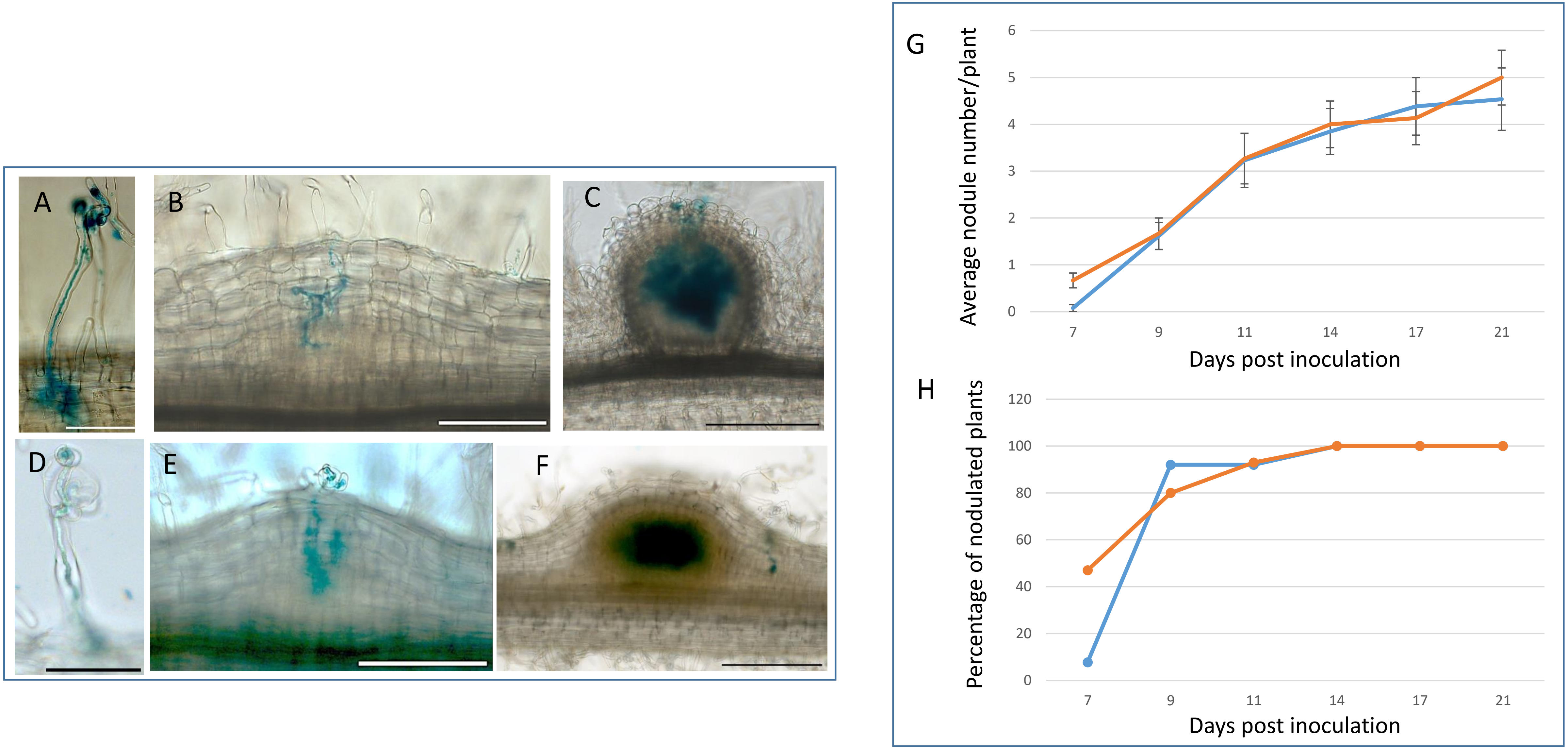


**Figure S15. Infection and kinetics of nodulation on *Mtsobir1-1* mutant and WT plants with *S. meliloti* 2011 in *in vitro* conditions.** A-F Photos illustrating the infection of the *Mtsobir1-1* mutant with *S. meliloti* 2011. A-C WT, D-F mutant. *S. meliloti* 2011 is coloured in blue by lacZ staining. Scale bars 50 µm (A, D), 100 µm (B, C, E, F). G, average nodule numbers per plant at different time points post inoculation. H, percentages of plants showing nodules at different time points post inoculation. Plants were grown on plates. Blue= *Mtsobir1-1* WT (n= 13), orange = *Mtsobir1-1* mutant (n=15).

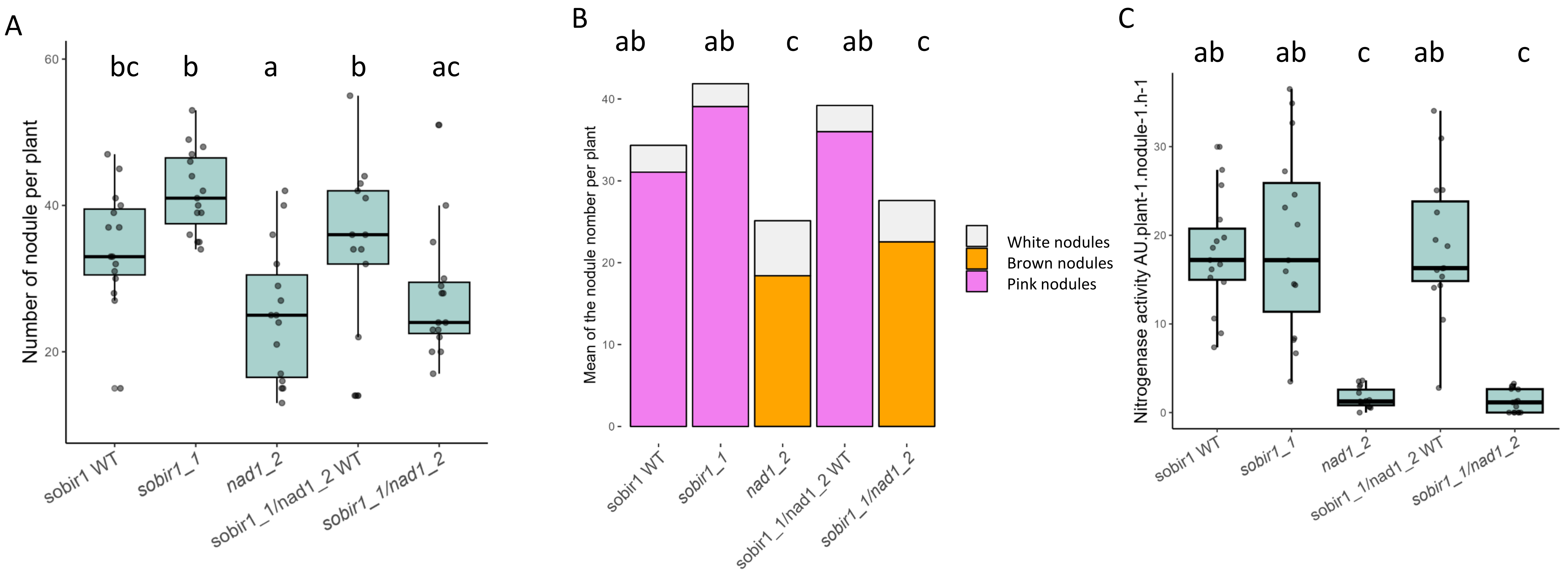


**Figure S16. Mt*SOBIR1* does not intervene in the activation of defence responses that are suppressed by *NAD1.*** The single *nad1-2* Mtsobir1-1 mutants as well as a double *sobir1-1/nad1-2* mutated WT control, were used for nodulation assays with WT *S. meliloti* strain 2011. A, The total number of nodules were scored at 21dpi. Lowercase letters indicate significant differences (ANOVA, Turkey, p<0.05). B, The nodule categories in different genotypes. White, pink and brown shading for white, pink and brown nodules, respectively. C, Nitrogenase activity of mutant and WT lines at 21dpi. 15 plants/genotype were used. Lowercase letters indicate significant differences. ANOVA, Turkey, p<0.05

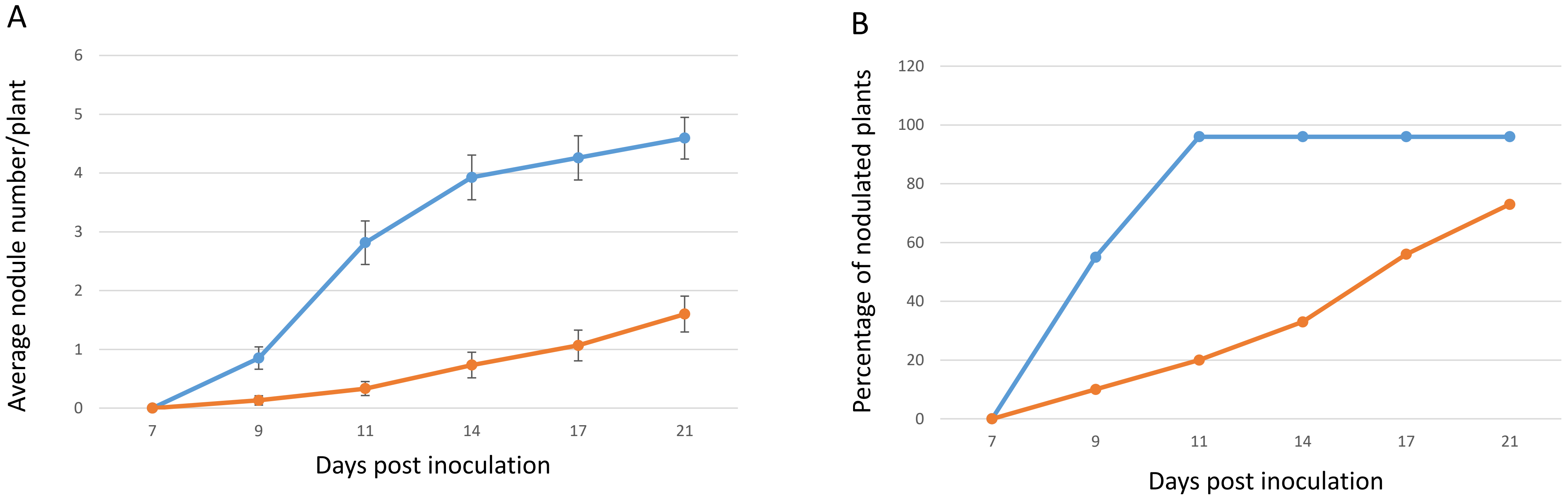


**Figure S17. Kinetics of nodulation on *Mtsobir1-1* mutant and WT plants with *S. medicae* WSM419 in *in vitro* conditions**
A,  average nodule numbers per plant at different time points post inoculation. B, percentages of plants showing nodules at different time points post inoculation. Data are from plate-grown plants with 2 independent repeats, blue= *Mtsobir1-1* WT (n= 27), orange = *Mtsobir1-1* mutant (n=30 ).

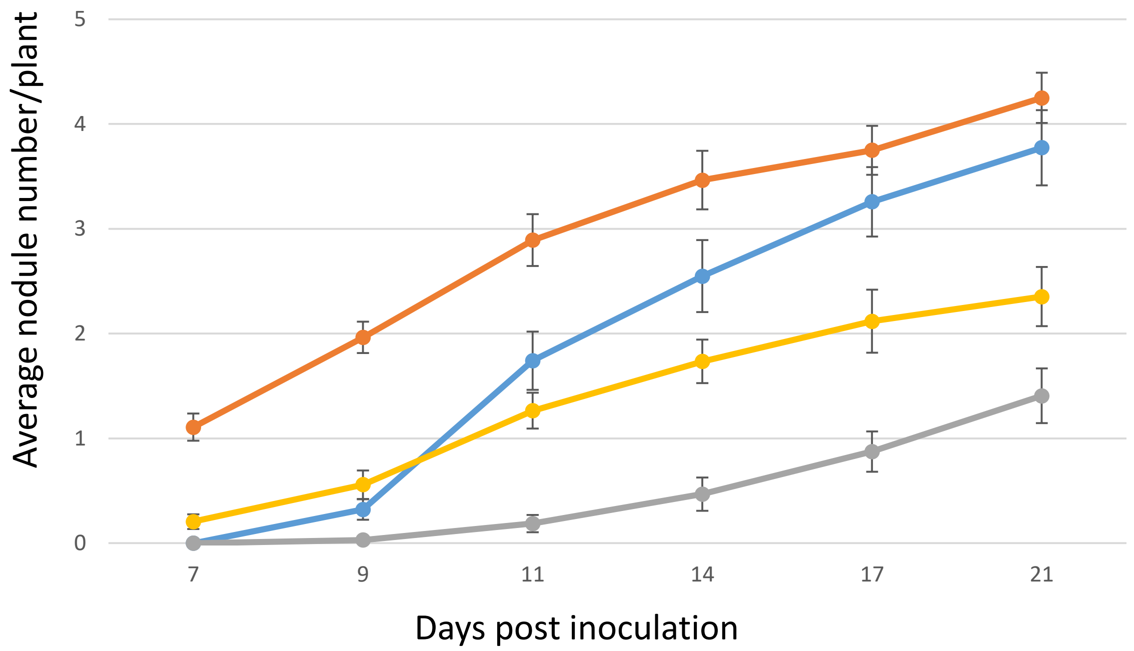


**Figure S18. Nodulation tests on *Mtsobir1-1* mutant and WT plants with *S. medicae* WSM419 in in vitro conditions with/without 100 mM AVG.** Average nodule numbers per plant at different time points post inoculation by *S. medicae* WSM419. Data are from plate-grown plants with 2 independent repeats; blue= WT without AVG (n= 31), orange = WT with AVG (n=28)**,** grey = mutant without AVG (n=32), yellow = mutant with AVG (n=34).

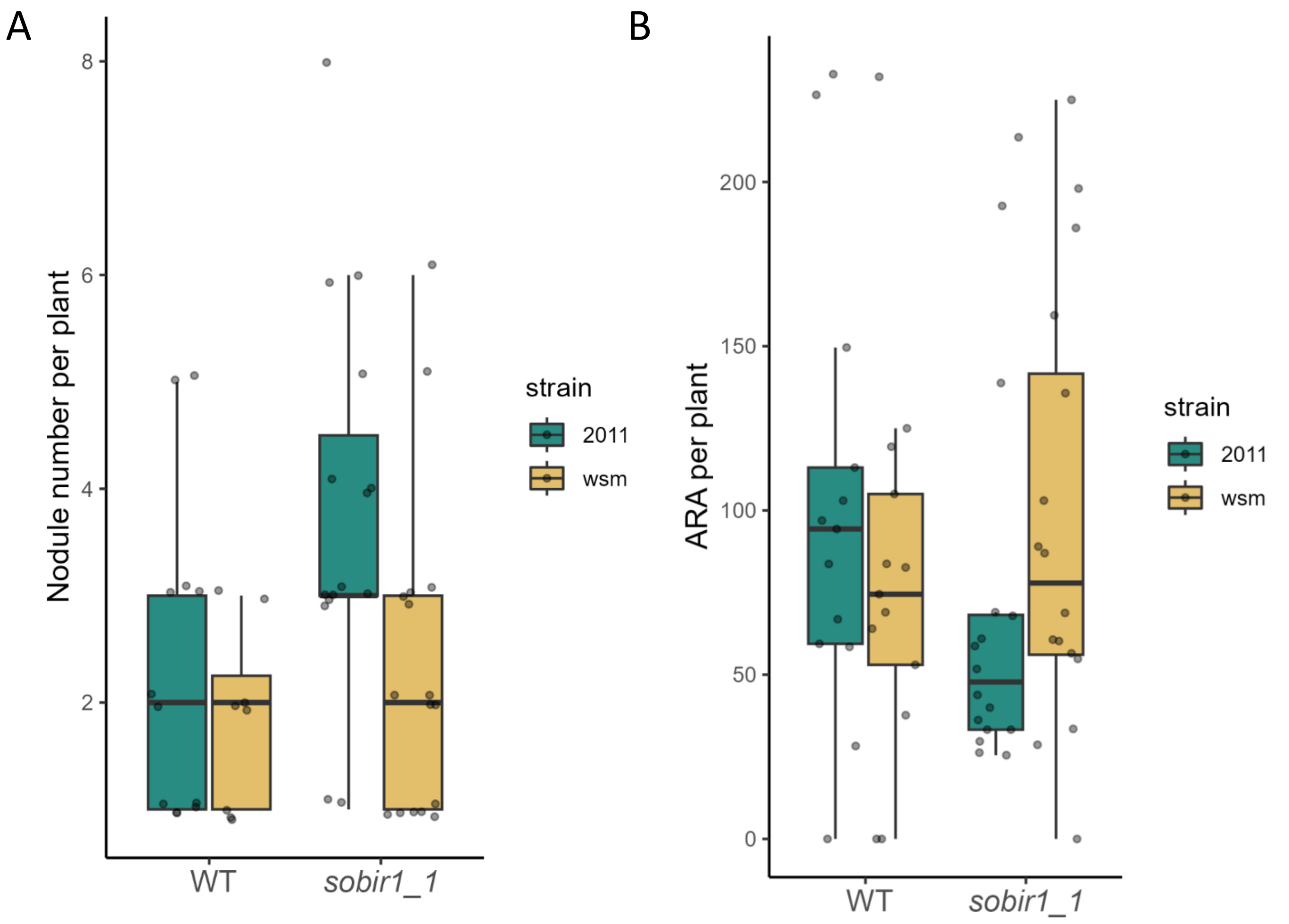


**Figure S19. Nodulation and nitrogenase activity of *Mtsobir1-1* plants inoculated by *S. meliloti 2011* or *S. medicae* WSM419**. Nodulation (A) and Acetylene Reduction Assays (ARA) (B) were assayed on *Mtsobir1-1* mutant and R108 (WT) plants nodulated by either *S. meliloti 2011* or *S. medicae* WSM419 in in vitro conditions at 21 dpi.

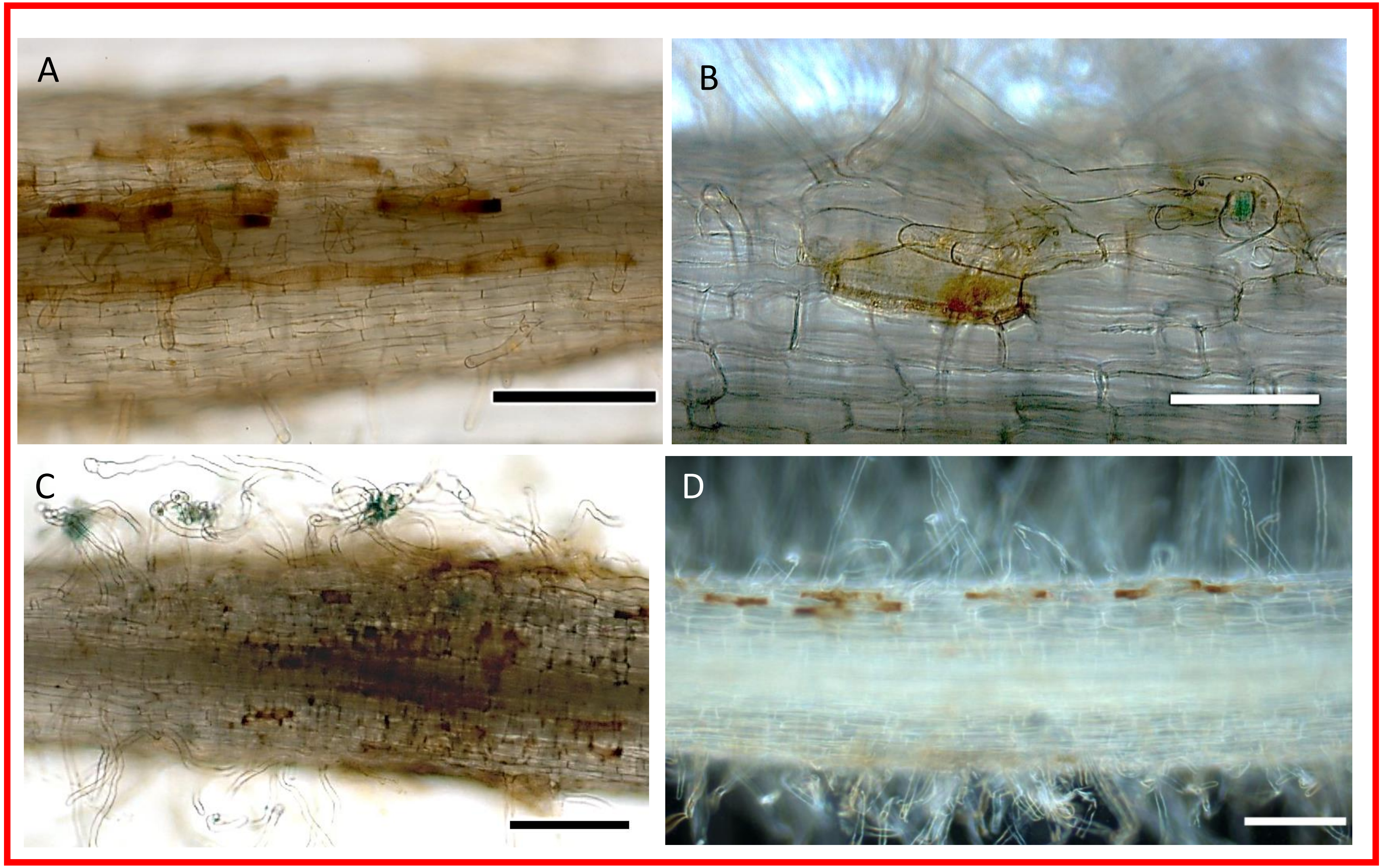


**Figure S20. Brown epidermal cells observed for *Mtsobir1-1* and *Mtsobir1-3* mutants when inoculated in *in vitro* conditions.** A-D, brown, necrotic cells observed in tube or plate-grown plants inoculated with either *S. meliloti* 2011 or *S. medicae* WSM419; A, *Mtsobir1-1/S. meliloti 2011;* B, *Mtsobir1-3/S. meliloti 2011;* C, *Mtsobir1-1/S. medicae* WSM419; D, *Mtsobir1-1/S. medicae* WSM419. Scale bars 100 µm (A, C, D), 50 µm (B)
